# Actin remodelling controls proteasome homeostasis upon stress

**DOI:** 10.1101/2022.02.09.479706

**Authors:** Thomas Williams, Roberta Cacioppo, Alexander Agrotis, Ailsa Black, Houjiang Zhou, Adrien Rousseau

## Abstract

When cells are stressed, bulk translation is often downregulated to reduce energy demands whilst stress-response proteins are simultaneously upregulated. 19S Regulatory Particle Assembly-Chaperones (RPACs) are selectively translated upon TORC1 inhibition to promote proteasome assembly and activity, maintaining cell viability. However, the molecular mechanism for such selective translational upregulation is unclear. Using yeast, we discover that remodelling of the actin cytoskeleton is important for RPAC translation following TORC1 inhibition. mRNA of the RPAC ADC17 travels along actin cables and is enriched at cortical actin patches under stress, dependent upon the early endocytic protein Ede1. *ede1*Δ cells failed to induce RPACs and proteasome assembly upon TORC1 inhibition. Conversely, artificially tethering ADC17 mRNA to cortical actin patches enhanced its translation upon stress. These findings suggest that actin dense structures such as cortical actin patches may serve as a translation platform for a subset of stress-induced mRNAs including regulators of proteasome homeostasis.

## Introduction

Cells require the right amount of each protein to be in the right place at the right time. This intimate balance between protein synthesis, folding, modification and degradation is known as protein homeostasis or proteostasis. Failure to maintain proteostasis causes a build-up of misfolded proteins, and potentially toxic aggregates, including those that cause neurodegenerative diseases^1–3^.

Misfolded, damaged, and short-lived proteins are degraded by one of two mechanisms: the autophagy-lysosome system, or the ubiquitin-proteasome system (UPS)^4,5^. In autophagy, autophagosomes are assembled around proteins or organelles to be degraded, then delivered to the lysosome^5^. With the UPS, proteins are tagged with ubiquitin conjugates which serve as a recognition signal for proteasomal degradation. The proteasome is a large, multiprotein complex comprising a ‘Core Particle’ (CP), containing the proteolytic activity of the proteasome, and one or two ‘Regulatory Particles’ (RP). The RP recognises ubiquitinated proteins and catalyses their unfolding and translocation into the CP, where they are degraded^6,7^.

Under optimal energy and nutrient conditions, protein synthesis exceeds degradation to achieve cell growth and proliferation. The TORC1 kinase complex (in yeast, mTORC1 in mammals) is active and promotes anabolic processes such as protein, nucleotide and lipid synthesis^8–10^. Concurrently, TORC1 restricts autophagy induction and proteasomal degradation^11–13^. When nutrients are limited, or certain stresses are applied to cells, TORC1 is inactivated^9,10^. TORC1 inactivation reduces bulk protein synthesis and increases protein degradation. Autophagy is de-repressed, while proteasomal degradation is increased. Enhanced protein degradation capacity allows cells to rapidly degrade misfolded and damaged proteins while simultaneously generating a pool of intracellular amino acids for stress-adaptive protein synthesis^14,15^. While the mechanism of TORC1 autophagy regulation has been well described, its regulation of proteasome function is much less clear. TORC1 inactivation in budding yeast leads to activation of the MAP Kinase Mpk1 (also known as Slt2, Erk5 in mammals)^13,16^. Activated Mpk1 is important to induce the translation of proteasome RP assembly chaperones (RPACs) such as Adc17 and Nas6, increasing RP assembly to produce more functional proteasomes^13^. CP assembly is also increased upon TORC1 inhibition with two CP assembly chaperones, Pba1 and Pba2, being induced. Unlike RPAC induction, CP assembly chaperone induction is Mpk1-independent, the underlying mechanism being unknown^13^. This increase in assembled proteasomes is transient and is followed by an overall reduction via proteaphagy^17^.

The mechanism of increased RPAC translation, while most protein synthesis is inhibited, is so far unknown. Here, we show that the endocytic protein Ede1 plays a key role in this process. Our findings indicate that actin remodelling is important for regulating the localisation and therefore the selective translation of mRNAs encoding stress-induced proteins such as RPACs.

## Results

### Identification of potential RPAC translation regulators

To better understand the mechanisms underlying selective RPAC translation, we established the FGH17 reporter system. The FGH17 reporter encodes two N-terminal <u>F</u>lag epitopes, a <u>G</u>FP and a C-terminal <u>H</u>A tag, under the control of the regulatory elements of the RPAC Adc<u>17</u> (Fig. 1a). FGH17-containing cells had low basal levels of FGH17, which strongly increased following rapamycin (TORC1 inhibitor) treatment, as for the endogenous RPAC Nas6 (Fig. 1a). We additionally compared the behaviour of endogenous ADC17 mRNA to FGH17 to confirm that they share similar translation regulation. Using RiboTag^18^, we observed that rapamycin increased the recruitment of both ADC17 mRNA and FGH17 mRNA to ribosomes for translation (Fig. 1b). This confirmed that FGH17 is a good reporter to interrogate how translation of mRNAs such as Adc17 are regulated upon stress. We next investigated the contribution of the UTRs to FGH17 translation regulation. Deletion of the 3’UTR had no effect on the regulation of the FGH17 reporter. In contrast, deletion of the 5’UTR abrogated FGH17 translation indicating that the 5’UTR of Adc17 contains the required translation regulation element(s) (Fig. 1c). Deletion of the 40 nucleotides upstream of the start codon (FGH17-40ntΔ) only slightly decreased translation of the FGH17 reporter, while deletion of the 70 nucleotides upstream of the start codon (FGH17-70ntΔ) prevented translation (Fig. 1d). This was not due to alteration of the Kozak sequence, as reintroducing ADC17 Kozak sequence to FGH17-70ntΔ (FGH17-70ntΔ+Kozak) was not enough to restore FGH17 expression (Extended data Fig. 1a). The 70-nucleotide region alone was not sufficient for FGH17 reporter expression (Extended data Fig. 1b). Comparing FGH17 with FGH17-70ntΔ by RiboTag, we observed that the deletion of this 70nt sequence prevented the recruitment of FGH17-70ntΔ mRNA to ribosomes upon rapamycin treatment (Fig. 1b, e) and decreased its stability by about 2-fold (Extended data Fig.1c). Deleting this 70nt region at the endogenous ADC17 locus with CRISPR/CAS9, we similarly observed abrogation of ADC17 expression (Fig. 1f). These findings indicated that the FGH17 reporter reflects the regulation of the endogenous *ADC17* gene.

**Fig. 1.**
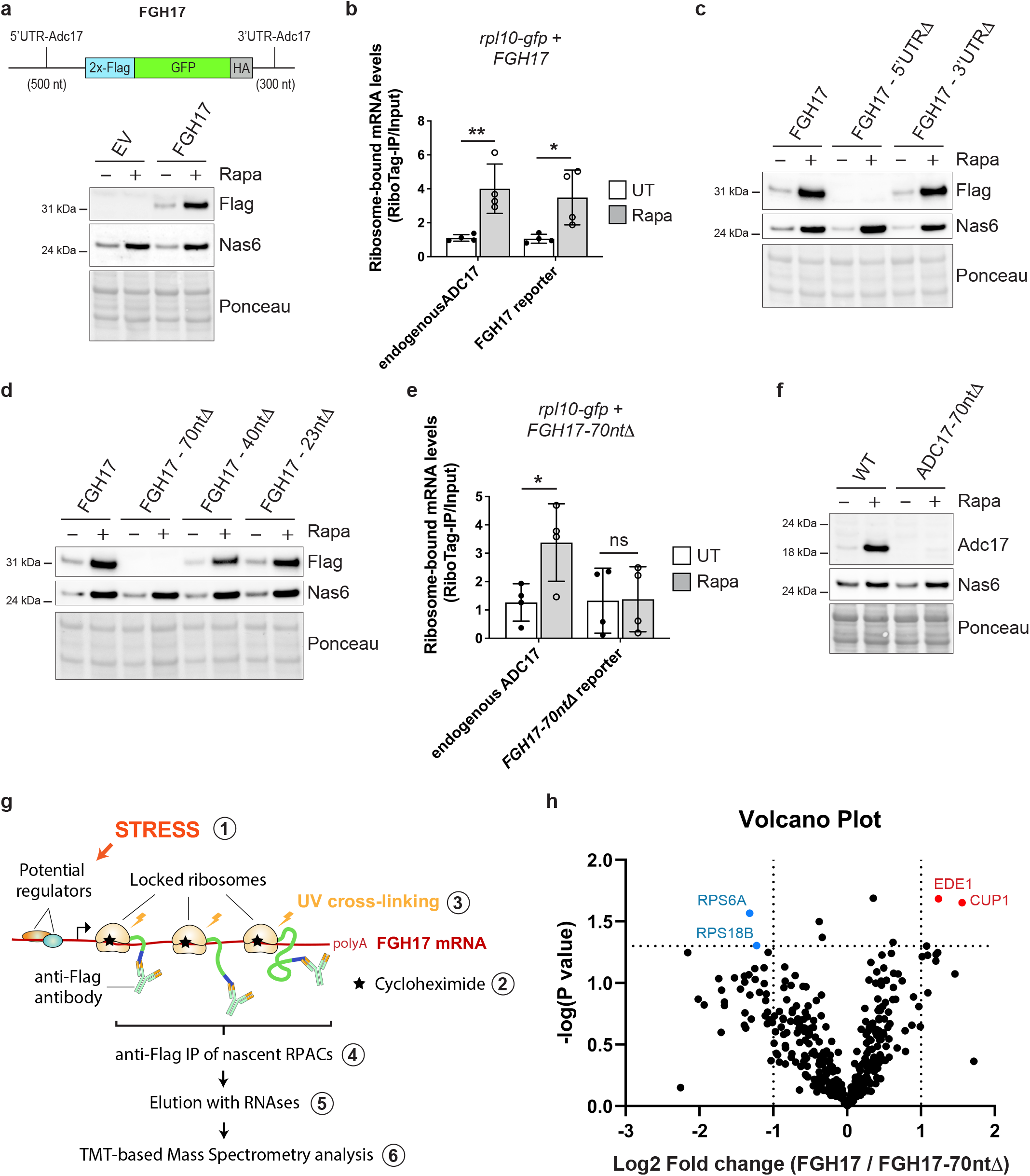
Identification of proteins interacting with translating RPAC reporter mRNAs. **a**, Cartoon depicting the RPAC translation reporter construct FGH17, consisting of tandem reporters expressed under control of the UTRs of Adc17 and western blot analysis of FGH17 expression in cells treated ± 200 nM rapamycin (Rapa) for 4 h. Ponceau S staining was used as a loading control. Empty vector (EV). **b,** mRNA levels of endogenous ADC17 and of FGH17 bound to ribosomes after 1.5 h rapamycin treatment compared to untreated cells. Analysis was performed by RiboTag IP followed by qRT-PCR and normalised to the housekeeping gene *ALG9.* Ribosome-bound mRNA corresponds to the level of RiboTag IP mRNA normalised to the level of Input mRNA. For each bar *n*=4 biologically independent experiments were performed. Statistical analysis was carried out using multiple unpaired t-test. \*\**P* ≤ 0.01; \**P* ≤ 0.05. **c–d**, Western blot analysis of WT and mutant FGH17 reporters (anti-Flag) and Nas6 in yeast cells treated ± 200 nM rapamycin (Rapa) for 4 h. Ponceau S staining was used as a loading control. **e,** mRNA levels of endogenous ADC17 and of FGH17-70ntΔ bound to ribosomes after 1.5 h rapamycin treatment compared to untreated cells. Analysis was performed as in **b**. For each bar *n*=4 biologically independent experiments were performed. \**P* ≤ 0.05; ns not significant. **f,** Western blot analysis of Adc17 and Nas6 expression in WT and ADC17-70ntΔ cells treated ± 200 nM rapamycin (Rapa) for 4 h. Ponceau S staining was used as a loading control. **g**, Cartoon depicting the proteomics experimental design. Step 1, Cells expressing WT or mutant FGH17 were treated ± 200 nM rapamycin for 1.5 h; Step 2, Ribosomes were locked on mRNAs by treating cells with 35 μM cycloheximide; Step 3, Cells were treated with 1.2J/cm^2^ UV to covalently crosslink proteins to RNA; Step 4, Translating FGH17 mRNAs where ribosomes have already synthesised at least one N-terminal Flag tag were immunoprecipitated; Step 5, Proteins bound to translating FGH17 mRNAs were recovered by RNase treatment before being subjected to tandem mass tag (TMT)-based quantitative proteomics (Step 6). **h**, Volcano plot showing the proteins which were differentially recovered from FGH17 and FGH17-70ntΔ immunoprecipitates. Each dot represents a protein. The red dots are proteins significantly more bound to FGH17 compared to FGH17-70ntΔ and blue dots are proteins significantly less bound to FGH17 compared to FGH17-70ntΔ. *n*=5 biologically independent samples per condition; *P* values were determined by multiple unpaired t-test. **a, c, d, f,** Data are representative of three independent biological replicates.

To discover new RPAC translation regulators, we identified RNA-binding proteins (RBPs) with increased recruitment to translating WT FGH17 mRNAs compared to non-translatable FGH17-70ntΔ mRNAs *in-vivo.* To this end, we treated yeast cells with rapamycin to stimulate FGH17 translation (Fig. 1g, step 1). Polysomes were stabilised by adding cycloheximide to the cells (Fig. 1g, step 2) and UV-crosslinking covalently linked the RNA and any bound proteins together (Fig. 1g, step 3). We next used anti-FLAG beads to immunoprecipitate translating FGH17-mRNA complexes where locked ribosomes had already synthesized one or both N-terminal FLAG tags (Fig. 1g, step 4). As FGH17-70ntΔ mRNAs are not translated, these samples should only immunoprecipitate proteins that bind non-specifically to the anti-FLAG beads. Based on the prediction that potential regulators of RPAC translation would be UV crosslinked to translating FGH17 mRNA (Fig. 1g), we used RNases to specifically elute these potential regulators (Fig. 1g, step 5). We identified proteins in the RNase elution by tandem mass tag (TMT)-based quantitative proteomics (Fig. 1g, step 6). Quantitative analysis (Fig. 1h) revealed that two proteins, Ede1 and Cup1, were enriched in the WT compared to FGH17-70ntΔ samples. In contrast, two proteins were significantly depleted in the WT samples (Rps6a and Rps18b) suggesting they could be translational repressors (Fig. 1h). Taken together, these results identify a region of ADC17 mRNA essential for translation and discover potential regulators of selective RPAC translation.

### Ede1 is important for RPAC induction following TORC1 inhibition

Defects in RPAC regulation sensitise yeast to rapamycin^13^. Therefore, to examine the involvement of identified proteins in RPAC regulation, we first tested knockout mutants for rapamycin sensitivity. Mutants of the ribosomal subunits Rps6a and Rps18b, which were less associated with translating FGH17 mRNA, were similarly sensitive to rapamycin as WT cells (Fig. 2a). Cup1 and Ede1 were found to be associated more with translating FGH17 mRNA and, while the *cup1Δ* mutant showed a similar level of sensitivity to rapamycin as WT cells, *ede1Δ* cells were highly sensitive (Fig. 2a).

**Fig. 2.**
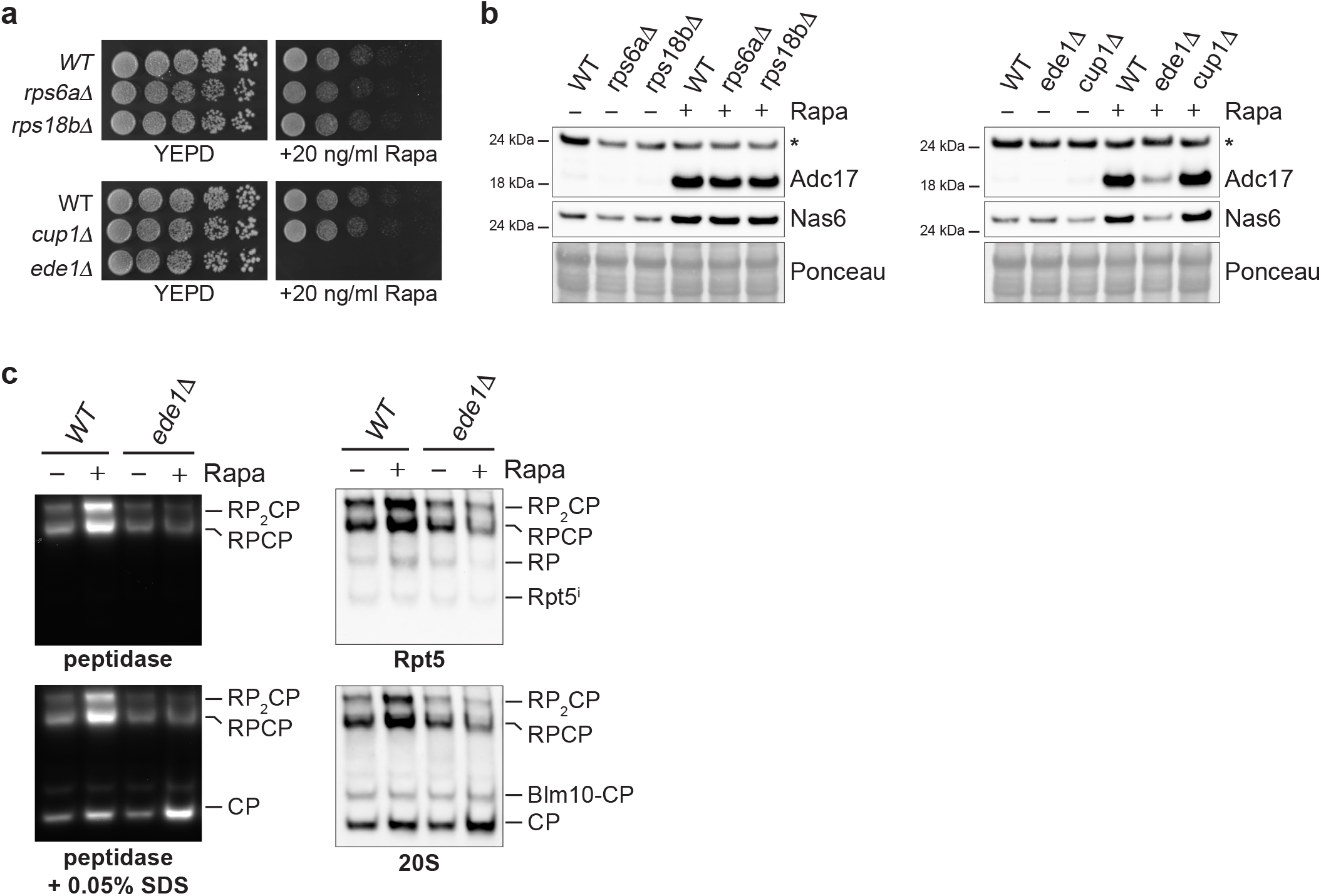
Ede1 regulates proteasome assembly upon TORC1 inhibition. **a**, Cells spotted in a fivefold dilution and grown for 3 days on plates ± 20 ng/ml rapamycin. **b**, Western blot analysis of RPACs in WT and deletion strains treated ± 200 nM rapamycin (Rapa) for 4 h. Ponceau S staining was used as a loading control. Asterisk indicates non-specific band. Data are representative of three independent biological replicates. **c**, Gradient Native polyacrylamide gel electrophoresis (PAGE) (3.8-5%) of yeast extracts from cells ± 200 nM rapamycin (Rapa) for 3 h, monitored by the fluorogenic substrate Suc-LLVY-AMC (left panel) and by immunoblots (right panel). Core particle (CP), single-capped (RPCP), double-capped (RP_2_CP) and Blm10-capped (Blm10-CP) proteasome complexes are indicated. Rpt5 and 20S antibodies recognise the RP and the CP, respectively. Data are representative of three independent biological replicates.

We next tested whether these mutants were defective in rapamycin-induced RPAC expression. Unlike *rps6aΔ, rps18bΔ* and *cup1Δ* cells, *ede1Δ* cells were severely impaired in both Adc17 and Nas6 induction following rapamycin treatment (Fig 2b), which, together with the rapamycin sensitivity, was rescued by reintroducing Ede1 (Extended data Fig. 2a, b). TORC1-mediated Rps6 phosphorylation was still inhibited following rapamycin treatment in *ede1Δ* cells, indicating that Ede1 acts downstream of TORC1 inhibition (Extended data Fig. 2c). A defect in RPAC induction following rapamycin treatment is expected to lead to a defect in increased proteasome assembly and activity^13^. Accordingly, *ede1Δ* cells failed to increase assembly and activity of the 26S proteasome, although there was an increase in 20S core particles (Fig. 2c). This defect is symptomatic of RP assembly defects and a hallmark of RPAC-deleted cells^13,19–21^. Ede1 is therefore necessary for enhanced proteasome assembly following rapamycin treatment, by increasing the amount of RPACs available.

### Ede1 interacts with ADC17 mRNA *in vivo* and regulates its translation

As Ede1 associates with translating FGH17 mRNA and is critical for proteasome homeostasis upon TORC1 inhibition, we predicted that Ede1 will be in contact with ADC17 mRNAs upon rapamycin treatment to play a role in their translation. To explore this possibility, PP7 stem loops were introduced into the endogenous Adc17 mRNA allowing it to be labelled with PP7 bacteriophage coat protein (PCP) fused to mKate2 in cells expressing GFPEnvy-tagged Ede1 (Fig. 3a). We tracked single molecules of labelled Adc17 mRNA and observed frequent contact of Ede1 and ADC17 mRNA, demonstrating that this interaction is occurring *in-vivo* (Fig. 3b and Supplementary Video 1,2). Around 29% of Adc17 mRNAs were associated to Ede1 under basal conditions and this significantly increased to about 40% following rapamycin treatment (Fig. 3c). This is consistent with a recent study that identified Ede1 as a potential RNA-binding protein^22^. To confirm that Ede1 is regulating ADC17 mRNA at the level of translation, we deployed the Suntag labelling method^23^. This method uses the multimerization of Gcn4 epitope (Suntag) which, when translated, is recognised by multiple single chain antibodies coupled to a fluorescent protein (scFv-mCherry), enabling quantitative visualization of the translation of individual mRNA molecules in living cells (Fig. 3d). We identified two populations of ADC17 mRNAs: those co-localising with Suntag signal which are translationally active (GFP and mCherry signal) (Fig. 3e, arrowheads 1 and 2) and those devoid of Suntag signal which are translationally inactive (GFP only) (Fig. 3e, arrowheads 3 and 4) (Supplementary video 3,4). On average, ~23% of ADC17 mRNAs were translationally active in untreated cells, increasing to ~40% in rapamycin-treated cells attesting that ADC17 mRNA translation is increased upon TORC1 inhibition (Fig. 3f). This increased translation of ADC17 mRNA was lost in *ede1Δ* cells treated with rapamycin, confirming the importance of Ede1 for Adc17 translation regulation (Fig. 3f). Taken together, these results show ADC17 mRNAs partly localise to Ede1 sites and demonstrate that Ede1 is critical for ADC17 mRNA translation upon TORC1 inhibition.

**Fig. 3.**
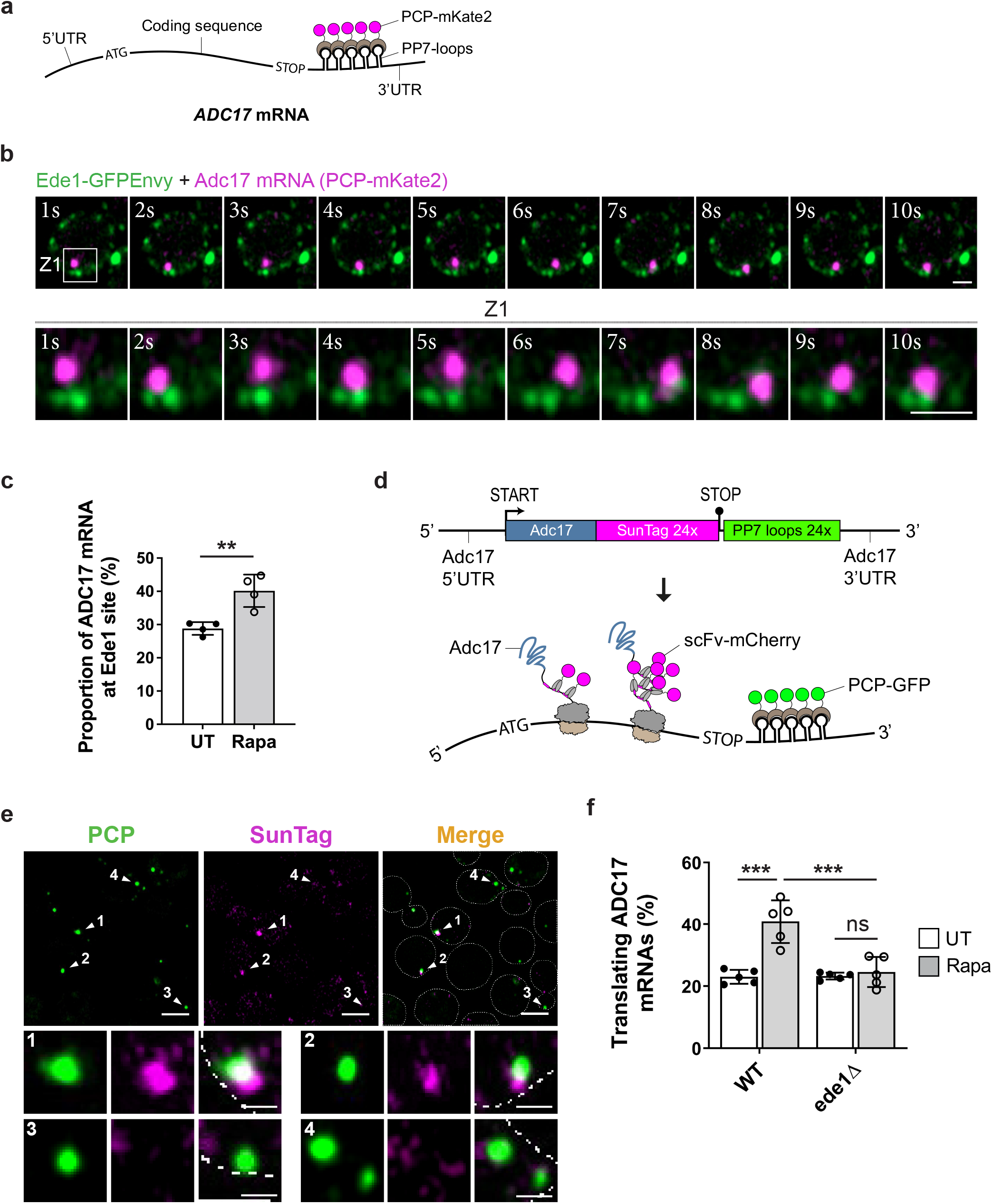
Ede1 controls ADC17 mRNA translation upon TORC1 inhibition. **a**, Cartoon depicting how single-molecule ADC17 mRNAs are labelled with PP7 bacteriophage coat protein (PCP) fused to mKate2 for fluorescence live-cell imaging. PP7 stem loops were introduced into the endogenous Adc17 mRNA allowing it to be selectively labelled with PCP-mKate2 in cells expressing Ede1-GFPEnvy. **b**, Montage from time-lapse imaging showing contacts between Ede1-GFPEnvy (green) and ADC17 mRNA (magenta). Scale bars, 1 μm. **c,** Frequency of ADC17 mRNAs (green) colocalising with Ede1-tdimer2 in cells grown for 3 h ± 200 nM rapamycin (Rapa). Untreated (UT). For each bar, *n*=4 biologically independent experiments with at least 500 ADC17 mRNAs for each condition. Statistical analysis was carried out using unpaired t-test.. \*\**P* ≤ 0.01. **d**, Schematic representation of ADC17-SunTag reporter mRNA used for single-molecule imaging of mRNA translation during stress. PCP-GFP labels Adc17 mRNA whereas scFv-mCherry labels translating Adc17 protein **e**, Representative microscopy images of yeast cells expressing ADC17-SunTag reporter mRNA. Translating ADC17 mRNAs are GFP (green)- and mCherry (magenta)-positive while non-translating ADC17 mRNAs are only positive for GFP. Translating ADC17 mRNAs are denoted by white arrowheads 1 and 2, while non-translating ADC17 mRNAs are denoted by white arrowheads 3 and 4. Higher magnification is shown at the bottom. Scale bars, 3 μm. **f**, Frequency of ADC17 mRNAs undergoing translation in WT and ede1Δ cells ± 200 nM rapamycin (Rapa) for 3 h using the Suntag labelling method. Untreated (UT). For each bar, *n*=5 biologically independent experiments with at least 500 ADC17 mRNAs for each condition. Statistical analysis was carried out using two-way ANOVA t-test (Tukey multiple comparison test). \*\*\**P* ≤ 0.001; ns, not significant.

### Ede1-associated actin regulatory proteins function in RPAC translation

Having established that Ede1 regulates RPAC translation, we investigated the underlying mechanism. Ede1 is an early coat protein involved in clathrin-mediated endocytosis (CME)^24^. To determine whether endocytosis is important for stress-mediated proteasome assembly, we tested if mutants of other non-essential proteins involved in CME mimicked the defects of *ede1Δ* cells. We initially screened mutants for increased rapamycin sensitivity, revealing 6 further proteins which were essential for growth on rapamycin (Fig. 4a). We examined whether these proteins were also involved in RPAC induction following rapamycin treatment. Chc1, Clc1, Pal1 and Vps1 were dispensable for induction of Adc17 and Nas6 after rapamycin treatment, however *sla1Δ* and *vrp1Δ* cells were severely impaired in RPAC induction (Fig. 4b). Like for *ede1Δ* cells, this was due to a translation defect, as observed using the Suntag labelling method (Fig. 4c). We next confirmed that *sla1Δ* and *vrp1Δ* cells had similar defects to *ede1Δ* in proteasome assembly and activity in response to rapamycin (Fig. 4d).

**Fig. 4.**
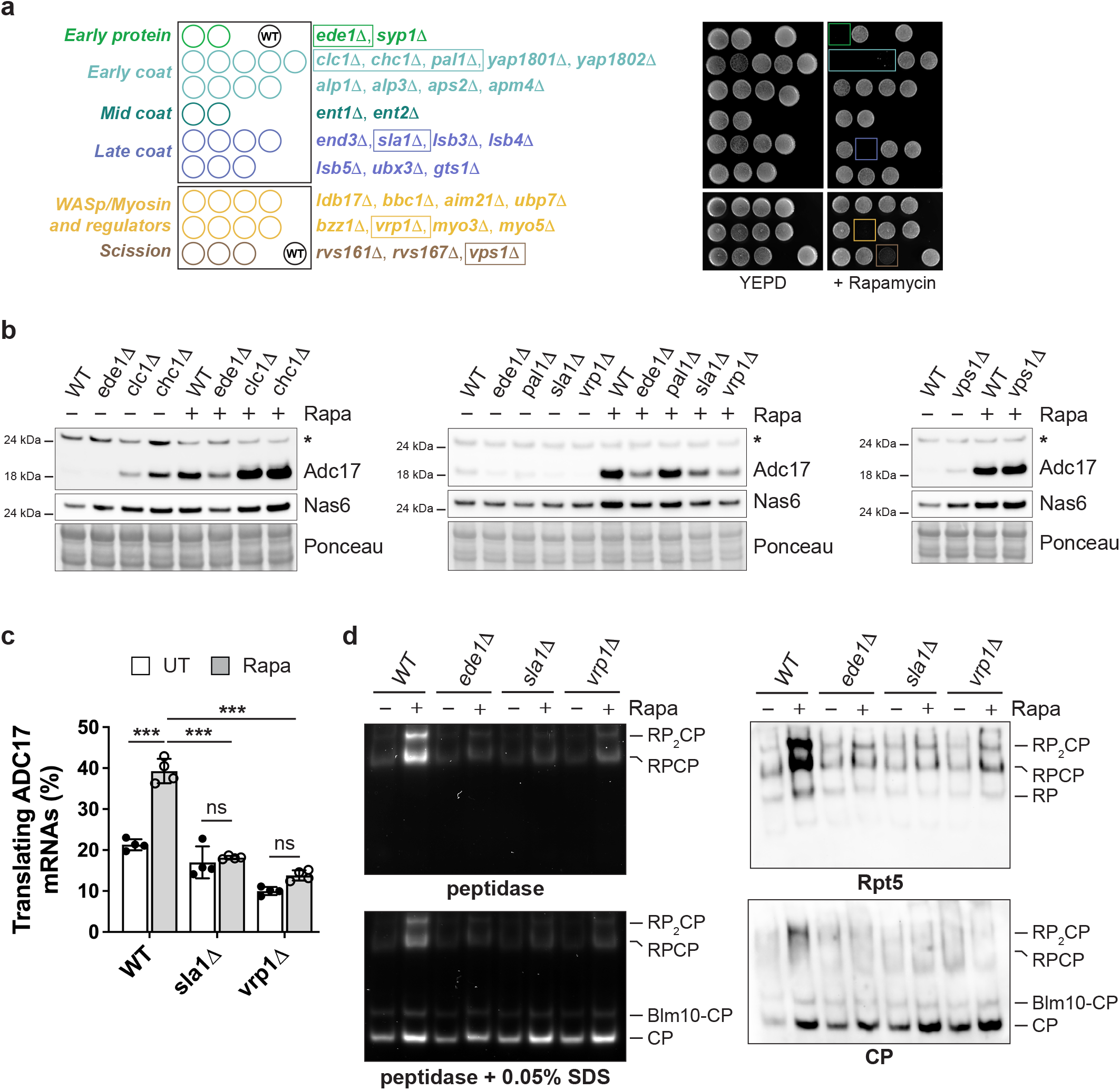
Ede1, Sla1 and Vrp1 are important for proteasome assembly and activity. **a**, Screen for rapamycin sensitivity with deletion strains covering all non-essential endocytic genes. Left panel, schematic representation of YEPD plates indicating the position of deletion strains. WT yeast was used as a control in the indicated positions. Right panel, yeast growth for 3 days on YEPD plate ± 20 ng/ml rapamycin from the indicated strains. Strains which failed to grow on rapamycin are indicated in coloured boxes in both the schematic and plate image. **b**, Western blot analysis of RPACs in WT and deletion strains treated ± 200 nM rapamycin (Rapa) for 4 h. Ponceau S staining was used as a loading control. Asterisk indicates non-specific band. **c**, Frequency of ADC17 mRNAs undergoing translation in WT, *sla1*Δ and *vrp1*Δ cells ± 200 nM rapamycin (Rapa) for 3 h using the Suntag labelling method. Untreated (UT). For each bar, *n*=4 biologically independent experiments with at least 500 ADC17 mRNAs for each condition. Statistical analysis was carried out using two-way ANOVA t-test (Tukey multiple comparison test). \*\*\**P* ≤ 0.001; ns, not significant. **d**, Gradient Native polyacrylamide gel electrophoresis (PAGE) (3.8-5%) of yeast extracts from cells ± 200 nM rapamycin (Rapa) for 3 h, monitored by the fluorogenic substrate Suc-LLVY–AMC (left panel) and by immunoblots (right panel). Core particle (CP), single-capped (RPCP), double-capped (RP2CP) and Blm10-capped (Blm10-CP) proteasome complexes are indicated. Rpt5 and 20S antibodies recognise the RP and the CP, respectively. **a, b, d,** Data are representative of three independent biological replicates.

### Actin remodelling affects Adc17 mRNA localisation

Ede1 is one of the first protein to be recruited at endocytic sites (Fig. 5a, step 1). Sla1 forms a heterodimeric complex with the actin nucleation promoting factor (NPF) Las17 which is recruited to endocytic patches via Sla1-Ede1 interaction (Fig. 5a, step 2). Vrp1 is recruited to the endocytic site by Las17 (Fig 5a, step 3), and contributes to the recruitment and the activation of Myo3 and Myo5 that have both NPF and motor activities (Fig. 5a, step 4). NPFs further recruit and activate the actin nucleator complex Arp2/3 to initiate actin nucleation (Fig. 5a, step 5)^25–27^. As Ede1, Sla1 and Vrp1 localise to and regulate cortical actin patches at the endocytic site, and ADC17 mRNA makes contacts with Ede1, Sla1 and Vrp1 (Fig. 3c and Extended data Fig. 3a, b), it seemed likely there might be a role for actin in Adc17 mRNA regulation. To test this, we fixed cells expressing endogenous PCP-GFP-labelled ADC17 mRNA and stained them with phalloidin to visualise actin. Yeast has two major actin structures: actin cables which are polarized linear bundles of parallel actin filaments extending along the long axis of cells and cortical actin patches which are dense dendritic networks of branched actin filaments localised at the plasma membrane^26^. ADC17 mRNA was mainly seen to localise on actin cables (~70%), with the remainder either on cortical actin patches (~26.5%) or not associated with any phalloidin staining (~3.5%) (Fig. 5b, c). We next sought to examine whether ADC17 mRNA travels along the actin cytoskeleton using live cell imaging. Tagged Abp1 and Abp140 were used to visualise cortical actin patches and cables, respectively. We observed that ADC17 mRNA is frequently transported along actin cables *in-vivo* (Fig. 5d and Supplementary Video 5,6), while its interaction with patches is more transient and dynamic, as previously observed for Ede1 (Extended data Fig. 3c and Supplementary Video 7,8). Overall, these data show that ADC17 mRNA travels along actin cables with the potential to either interact with cortical actin patches or be released from actin.

**Fig. 5.**
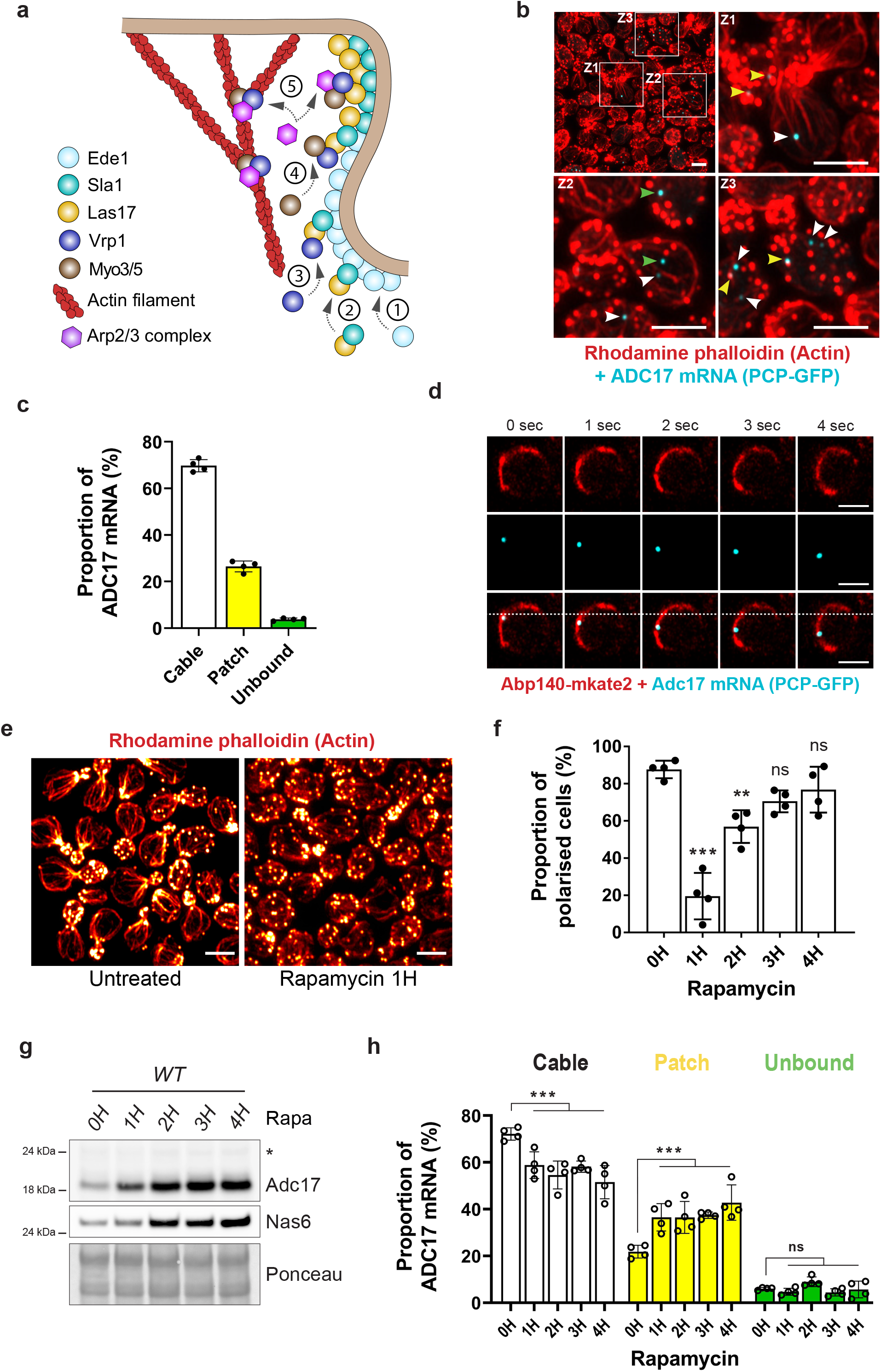
ADC17 mRNA travels along actin cables and relocalises to patches upon stress. **a,** Cartoon depicting the role of Ede1, Sla1 and Vrp1. Step 1, Ede1 is recruited to nascent endocytic sites. Step 2, Sla1 binds the nucleation promoting factor (NPF) Las17 and recruits it to endocytic sites, aided by the presence of Ede1. Step 3, Vrp1 is recruited to the endocytic site by Las17. Step 4, Vrp1 helps recruiting Myo3 and Myo5 that have both NPF and motor activities. Step 5, The NPFs Las17, Myo3 and Myo5 recruit and activate the actin nucleator complex Arp2/3 for cortical actin patch synthesis. **b**, Representative microscopy images (maximum intensity Z-projection) of yeast cells containing the PCP-GFP-labelled ADC17 mRNA (cyan) and stained with Rhodamine phalloidin to visualise actin (red). Z1, Z2 and Z3 areas are shown at higher magnifications. White, yellow and green arrowheads indicate ADC17 mRNAs bound to actin cable, cortical actin patch and not associated to actin, respectively. Scale bars, 3 μm. **c**, Frequency of ADC17 mRNAs bound to actin cable, cortical actin patch and not associated to actin structures. For each bar, *n*=4 biologically independent experiments with at least 200 ADC17 mRNAs for each condition. Cortical actin patches are defined as circular punctae with a diameter of 150–750 nm. Actin cables are defined as linear structures with a length > 1 μM. **d**, Representative microscopy images from time-lapse imaging showing the transport of ADC17 mRNA along an actin cable. Actin cables and ADC17 mRNA are shown in red (Abp140-mKate2) and cyan (PCP-GFP), respectively. Scale bars, 3 μm. **e**, Representative microscopy images (maximum intensity Z-projection) of yeast cells treated ± 200 nM rapamycin for 1 h and stained with Rhodamine phalloidin to visualise actin (hot red LUT). Scale bars, 3 μm. **f**, Frequency of polarised cells following rapamycin (200 nM) treatment for the indicated time. For each bar, *n*=4 biologically independent experiments with at least 250 cells per condition. **g**, Western blot analysis of RPACs in WT cells treated ± 200 nM rapamycin (Rapa) for the indicated time. Ponceau S staining was used as a loading control. Data are representative of three independent biological replicates. **h**, Frequency of ADC17 mRNA bound to actin cable, cortical actin patch or not associated to actin in WT cells treated ± 200 nM rapamycin for the indicated time. For each bar, *n*=4 biologically independent experiments with at least 200 ADC17 mRNAs for each condition. **f**, **h**, Statistical analysis was carried out using one-way ANOVA t-test (Dunnett multiple comparison test). \*\**P* ≤ 0.01; \*\*\**P* ≤ 0.001; ns, not significant.

The distribution of cortical actin patches and cables is polarized in budding yeast with cortical actin patches being found almost exclusively in the bud, and cables being aligned longitudinally from the mother cell into the bud^26^. It has been reported that rapamycin depolarises the actin cytoskeleton^16^, and we have shown that ADC17 mRNA is largely localised to actin structures (Fig. 5b-d). It is possible, therefore, that actin depolarisation is a key step in RPAC induction. We first monitored the kinetics of actin depolarisation after rapamycin treatment. Budding cells containing >6 cortical actin patches in the larger mother cell were considered to have a depolarized actin cytoskeleton, as previously described^28,29^. Rapamycin rapidly induced actin depolarisation, peaking at 1H and returning to near-normal levels at 4H (Fig. 5e, f). RPAC induction coincides with actin depolarisation indicating that actin remodelling may relocate RPAC mRNAs to trigger their translation (Fig. 5g). To test this possibility, we tracked ADC17 mRNA in cells stained for actin. Rapamycin treatment induced a shift of ADC17 mRNA localisation from actin cables to patches, from 1 h (1.7-fold increase) onward compared to untreated cells (Fig. 5h). These results show actin depolarisation upon TORC1 inhibition relocates Adc17 mRNAs from actin cables to cortical actin patches which could be important for its selective translation.

### Actin depolarisation by Latrunculin B induces proteasome assembly and activity

We next tested whether an alternative means of selectively disrupting actin cables induced RPAC translation. Cells were therefore treated with 25 μM Latrunculin B, which at this concentration only disrupts actin cables^30^, as observed upon rapamycin treatment. Lat-B treatment completely abolished actin cables and thereby relocated ADC17 mRNA to either cortical actin patches or to an unbound state (Fig. 6a, b). Analysing the pathway regulating RPAC levels, we showed that Lat-B activates Mpk1 kinase (Fig. 6c). RPAC expression is induced straight after Mpk1 activation, as previously reported for rapamycin^13^ (Fig. 6c). This was further confirmed using genetic disruption of actin cables in the temperature-sensitive mutant *act1-101* (Extended data Fig. 4). As the RPAC level mirrors the level of proteasome assembly, we monitored the impact of Lat-B on proteasome activity. In-gel peptidase assays showed that, as for rapamycin, Lat-B is a potent inducer of proteasome assembly and activity (Fig. 6d), attesting that actin remodelling regulates proteasome homeostasis.

**Fig. 6.**
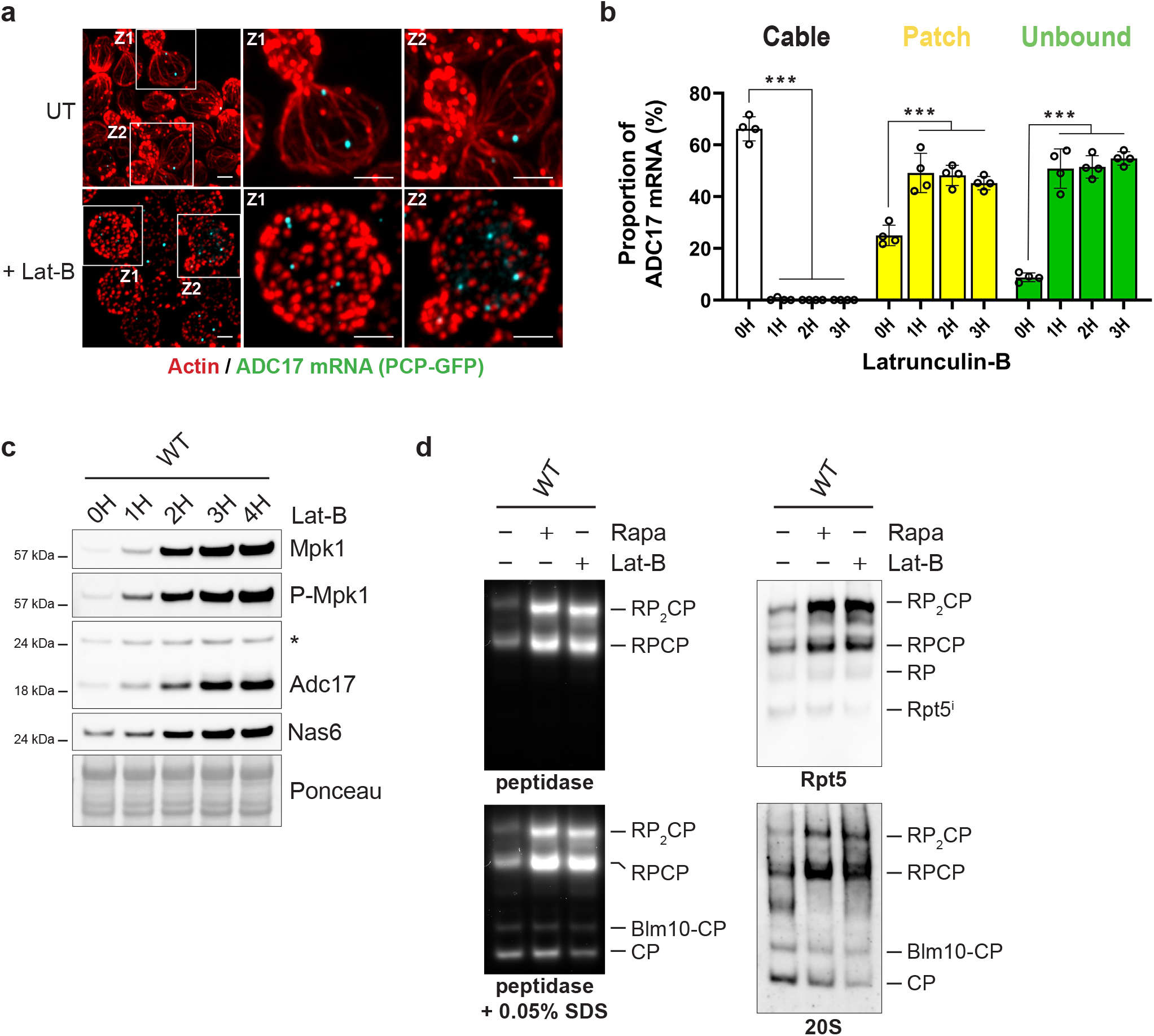
Latrunculin-B induces RPACs and proteasome assembly. **a,** Representative microscopy images (maximum intensity Z-projection) of yeast cells containing the PCP-GFP-labelled ADC17 mRNA (cyan), ± 25 μM Latrunculin-B (Lat-B) for 1 h and stained with Rhodamine phalloidin to visualise actin (red). Z1 and Z2 areas are shown at higher magnifications. Scale bars, 2 μm. **b**, Frequency of ADC17 mRNA bound to actin cable, cortical actin patch or not associated to actin in WT cells treated ± 25 μM Latrunculin-B for the indicated time. For each bar, *n*=4 biologically independent experiments with at least 200 ADC17 mRNAs for each condition. Statistical analysis was carried out using one-way ANOVA t-test (Dunnett multiple comparison test). \*\*\**P* ≤ 0.001. **c**, Western blot analysis of RPACs and Mpk1 kinase in WT cells treated ± 25 μM Latrunculin-B (Lat-B) for the indicated time. Ponceau S staining was used as a loading control. **d**, Gradient Native polyacrylamide gel electrophoresis (PAGE) (3.8-5%) of yeast extracts from cells ± 200 nM rapamycin (Rapa) or 25 μM Latrunculin-B (Lat-B) for 3 h, monitored by the fluorogenic substrate Suc-LLVY–AMC (left panel) and by immunoblots (right panel). Core particle (CP), single-capped (RPCP), doublecapped (RP2CP) and Blm10-capped (Blm10-CP) proteasome complexes are indicated. Rpt5 and 20S antibodies recognise the RP and the CP, respectively. **c**, **d**, Data are representative of three independent biological replicates.

### Ede1-mediated stabilisation of Adc17 mRNA at cortical actin patches enhances its translation following TORC1 inhibition

Having found that Lat-B induces RPAC expression, we sought to determine whether Ede1 is required for this process, as for rapamycin. Despite actin becoming depolarised after Lat-B treatment (Fig. 7a), RPACs were not induced in *ede1Δ* cells (Fig. 7b). This result shows that Ede1 plays a role downstream of actin depolarisation, possibly by stabilising Adc17 mRNAs at cortical actin patches. While rapamycin treatment induced a shift of ADC17 mRNA localisation from actin cables to cortical actin patches in WT cells, this re-localisation was lost in *ede1Δ* cells (Fig. 7c). This confirms Ede1 helps stabilise Adc17 mRNA at cortical actin patches following TORC1 inhibition.

**Fig. 7.**
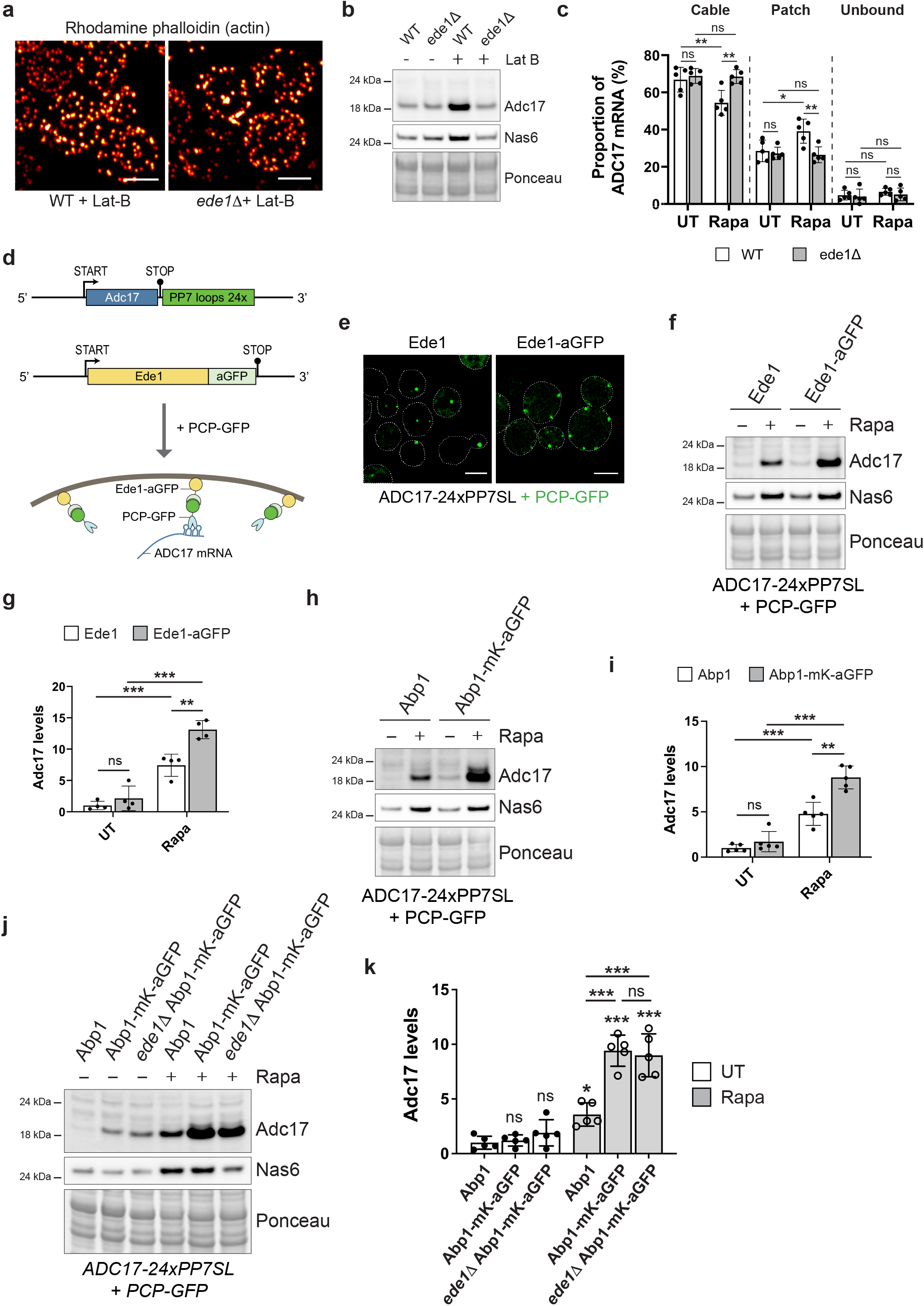
Tethering of ADC17 mRNA to actin patches enhances its translation upon stress. **a,** Representative microscopy images (maximum intensity Z-projection) of WT and *ede1*Δ cells treated with 25 μM Latrunculin B (Lat-B) for 1 h and stained with Rhodamine phalloidin to visualise actin (hot red LUT). Scale bars, 3 μm. **b**, Western blot analysis of RPACs in WT and *ede1*Δ cells treated ± 25 μM Latrunculin-B (Lat-B) for 3 h. Ponceau S staining was used as a loading control. Data are representative of three independent biological replicates. **c**, Frequency of ADC17 mRNA bound to actin cable, cortical actin patch or not associated to actin in WT and *ede1*Δ cells treated ± 200 nM rapamycin (Rapa) for 1 h. Untreated (UT). For each bar, *n*=5 biologically independent experiments with at least 200 ADC17 mRNAs for each condition. **d**, Schematic representation of the system used to artificially tether ADC17 mRNA to Ede1. PCP: PP7 bacteriophage coat protein, aGFP: nanobody against GFP. **e**, Representative microscopy images of yeast cells containing PCP-GFP-labelled ADC17 mRNA (green) and expressing either WT Ede1 or Ede1 tagged with a nanobody against GFP (Ede1-aGFP). Scale bars, 3 μm. **f**, Western blot analysis of RPACs in cells containing PCP-GFP-labelled ADC17 mRNA and expressing either WT Ede1 or Ede1 tagged with a nanobody against GFP (Ede1-aGFP), treated ± 200 nM rapamycin (Rapa) for 4 h. Ponceau S staining was used as a loading control. Data are representative of four independent biological replicates. **g**, Quantification of Adc17 protein level from experiments represented in **f**. Untreated (UT). For each bar, *n*=4 biologically independent experiments. **h**, Western blot analysis of RPACs in cells containing PCP-GFP-labelled ADC17 mRNA and expressing either WT Abp1 (Abp1) or Abp1-mKate2 tagged with a nanobody against GFP (Abp1-mK-aGFP), treated ± 200 nM rapamycin (Rapa) for 4 h. Ponceau S staining was used as a loading control. Data are representative of five independent biological replicates. **i**, Quantification of Adc17 protein level from experiments represented in **h**. Untreated (UT). For each bar, *n*=5 biologically independent experiments. **j**, Western blot analysis of RPACs in cells containing PCP-GFP-labelled ADC17 mRNA and expressing; WT Abp1 (Abp1), Abp1-mKate2 tagged with a nanobody against GFP (Abp1-mK-aGFP), or Abp1-mKate2 tagged with a nanobody against GFP with *EDE1* deleted (*ede1*Δ Abp1-mK-aGFP), treated ± 200 nM rapamycin (Rapa) for 4 h. Ponceau S staining was used as a loading control. Data are representative of five independent biological replicates. **k**, Quantification of Adc17 protein level from experiments represented in **j**. Untreated (UT). For each bar, *n*=5 biologically independent experiments. **c, g, i, k,** Statistical analysis was carried out using two-way ANOVA t-test (Tukey multiple comparison test). \**P*≤0.05; \*\**P*≤0.01; \*\*\**P*≤ 0.001; ns, not significant.

If stabilisation of Adc17 mRNAs at Ede1 sites is important for RPAC translation, artificially targeting Adc17 mRNA to this location may impact its translation levels upon TORC1 inhibition. Therefore, we set out to establish a targeting system in which Ede1 is fused to a nanobody recognising GFP (Ede1-aGFP) that will recruit the PCP-GFP proteins and thence the ADC17 mRNAs containing the PP7 stem loops (Fig. 7d). We first confirmed that PCP-GFP proteins are tethered to the plasma membrane where Ede1 is localised (Fig. 7e). Using a doubly tagged version of Ede1 (Ede1-tdimer2-aGFP), we also demonstrated that the PCP-GFP proteins are colocalising with Ede1 proteins (Extended data Fig. 5a, b). In this system, Ede1 sites are decorated with PCP-GFP and consequently all PCP-GFP dots do not correspond to one ADC17 mRNA molecule. Because of this, we validated that ADC17 mRNAs are indeed tethered to Ede1 sites using a doubly tagged version of ADC17 mRNA (Adc17-PP7SL-MS2SL). We observed that ADC17 mRNAs are robustly tethered to Ede1-aGFP/PCP-GFP sites indicating that our mRNA targeting system is efficient (Extended data Fig. 5c, d). Moreover, the tethering of ADC17 mRNA to Ede1 sites had no impact on its stability (Extended data Fig. 5e). We thus used this system to monitor the impact of artificially tethering ADC17 mRNAs to Ede1 sites on their translation level. We observed that Adc17 induction upon rapamycin treatment was increased around two-fold when tethered to Ede1-aGFP/PCP-GFP compared to untethered mRNAs, indicating recruitment of ADC17 mRNA to Ede1 sites is important for its translation regulation (Fig. 7f, g). Because Nas6 mRNA does not possess the PP7 stem-loops, its induction by rapamycin was unaffected by Ede1-aGFP (Fig. 7f).

As Ede1 localises to cortical actin patches, its main function in regulating RPAC translation could be to stabilise RPAC mRNAs at these sites. Therefore, we artificially tethered ADC17 mRNA to the cortical patch marker Abp1 and monitored RPAC levels. We observed that Adc17 induction upon rapamycin treatment was increased by 1.84-fold when tethered to Abp1 (Abp1-mK-aGFP) compared to untethered mRNAs (Abp1) which is similar to that of Ede1 targeting (Fig.7h, i). When ADC17 mRNA was targeted to cortical actin patches in *ede1Δ* cells, Adc17 induction upon rapamycin was restored to WT levels, indicating that an important function of Ede1 is to recruit ADC17 mRNA to cortical actin patches (Fig. 7j, k). Taken together, these results show that Ede1-mediated recruitment of ADC17 mRNAs at cortical actin patches following rapamycin treatment is important for stimulating Adc17 translation.

## DISCUSSION

Proteostasis maintenance under adverse conditions needs coordinated regulation of protein synthesis and degradation to provide adequate levels of required proteins, while clearing misfolded and damaged proteins^1,2,31^. Translational control is an integral part of the proteostasis network which allows fast and efficient proteome reprograming ^32,33^. Most stresses have been reported to repress overall protein synthesis while inducing selective translation of subsets of mRNAs that are essential for stress recovery. Here, we have discovered that remodelling of the actin cytoskeleton is critical to induce selective translation of RPAC mRNAs upon stress to adapt proteasome assembly to the cell requirements.

Local mRNA translation has been described as important for various processes, including development, cell migration and neuron function^35–37^. Yeast mRNAs have been reported to localise to the bud-tip, the ER, mitochondria and cortical actin patches, where they have either been shown, or are predicted, to be locally translated^38–41^. The most well-characterised of these is the ASH1 mRNA, which is transported along the actin cytoskeleton in a translationally repressed state to the bud tip, where it is anchored, and translation activated. It is possible a similar mechanism is responsible for RPAC induction. In this scenario, RPAC mRNA is transported along actin cables in a translationally repressed state. When actin cables are disrupted by stress, ADC17 mRNA relocalises to Ede1 sites and translation inhibition is relieved. Likewise, in human cells, mRNAs have been reported to localise to the ER, mitochondria, distal parts of neurons and actin dense structures such as focal adhesions, which are somewhat akin to cortical actin patches^42–45^. The localisation of certain mRNAs has been shown to be dependent on F-actin, while other mRNAs are transported along microtubules^41,48–50^. Here we have shown that Adc17 RPAC mRNA is transported on actin cables and interacts with cortical actin patches. Upon rapamycin and Latrunculin-B treatment, which respectively weaken and remove actin cables, the mRNA is stabilised at cortical actin patches. This relocalisation of RPAC mRNA allows increased RPAC translation and ultimately proteasome assembly in the presence of Ede1, Sla1 and Vrp1.

As cortical actin patches have a higher density of F-actin and are less sensitive to stress than actin cables, they may serve directly as a translation platform or indirectly by recruiting mRNA to a translationally active cellular compartment, helping to translate stress-induced proteins such as RPACs. In agreement with this possibility, it has been recently reported that Ede1 foci are surrounded by fenestrated ER containing membrane-associated ribosomes^49^. Recent findings showed that the ER has a far more diverse role in mRNA translation than expected, with ER-bound ribosomes functioning in the synthesis of both cytosolic and ER-targeted proteins^42^. These observations have been reported in diverse organisms and using different methodologies, suggesting that the ER is a favourable environment for translation. Furthermore, ER-localized mRNA translation is less inhibited by stress than their cytosolic counterparts, suggesting the ER represents a protective environment for translation upon stress, allowing the cell to synthesise a specific set of proteins under these conditions^42^. This would support a model in which the role of Ede1 in RPACs translation upon stress may be to recruit their mRNAs to cortical actin patches, so they are near ER-associated ribosomes for translation. As the ER has been reported to be an important site for 20S proteasome assembly, it would be possible to imagine that proteasome component are co-translationally assembled at the surface of ER membrane^50^.

In mammals, various actin dense structures have been described including actin meshes, foci, bundles and patches^51–53^. In the brain, reorganisation of actin dense structures allows a rapid remodelling of dendritic shafts and spines which is critical for neuronal plasticity, learning and memory^54,55^. The major architectural components of dendritic spines are long- and short-branched actin filaments formed by the Arp2/3 complex and NPFs, which are similar to that of cortical actin patches. Acute inhibition of the proteasome blocks activity-induced spine outgrowth, indicating that both proteasome and actin cytoskeleton are required for dendritic spine outgrowth and activity^56,57^. It has also been reported that the 26S proteasome is enriched at dendritic spines upon synaptic stimulation, with ~50% of the proteasomes being associated with the actin cytoskeleton^58^. As dendritic spines are translationally active, it is possible that mRNA encoding proteasome components can be transported to these structures and locally translated, although this remains to be studied.

Multiple stresses impact upon the actin cytoskeleton, leading it to be proposed as a biosensor for detecting stress^59^. In this work, we showed that rapamycin and Lat-B treatment perturb the actin cytoskeleton and re-localise a stress-responsive mRNA to cortical actin patches. This could potentially allow local translation of the mRNA throughout the mother cell, helping it to cope with the stress before resuming cell growth. While it is becoming clear that local and selective translation is crucially important in regulating cellular functions, less is known about how it is regulated under stress. This work illustrates that actin remodelling controls mRNA localisation and translation under stress, helping to adapt the proteome to environmental and physiological challenges. As perturbation of the actin cytoskeleton has been associated with various human diseases such as cancer and autoimmunity, it will be important to better understand the impact of actin cytoskeleton remodelling in controlling selective translation under pathophysiological conditions.

## Acknowledgements

We thank A. Bertolotti for the gift of Adc17 antibody. We thank the MRC PPU reagents and Services including Cloning team and DNA sequencing services, MRC-PPU Mass Spectrometry and the Dundee Imaging facility. We thank K. Labib and D. Alessi for reading and discussing the manuscript. This research was supported by the Medical Research Council (grant number MC_UU_00018/8 to AR).

## Author contributions

A.R. and T.W. designed experiments. A.R., T.W. wrote the manuscript. A.R., T.W., R.C., A.A., A.B., and H. Z performed experiments and analysed data. All authors edited the manuscript.

## Methods

### Yeast strains, plasmids and growth conditions

Yeast strains and plasmids used in this study are listed in Supplementary Table 1. All plasmids during this study were made with In-Fusion HD Cloning Plus (Takara, 638909). All yeast strains are isogenic to BY4741. All gene deletions were created using the PCR-based integration system^60^ and verified by PCR analysis. Cells were cultured in YEPD (yeast extract peptone) (2% glucose) or SC (synthetic medium) (2% glucose) lacking appropriate amino acids for selection. For yeast treatment, cells were grown on plates overnight at 30°C, resuspended to OD_600nm_ 0.2 in YEPD and grown at 30°C until they reached OD_600nm_ 0.5-0.8 to ensure exponential growth. Cells were diluted back to OD_600nm_ 0.2 before being treated with either 200 nM Rapamycin (LC Laboratories, R-5000), 25 μM Latrunculin-B (Abcam, ab144291), or shifted to non-permissive temperature for the indicated time. To assess growth using drop assays, yeast strains were adjusted to OD_600nm_ 0.2 from freshly streaked yeast and 5 μl of 1/5 serial dilutions spotted on YEPD plates ± 20 ng/ml rapamycin. Plates were imaged after 3 days at 30°C using the Chemidoc Touch imaging system (BioRad) on colorimetric setting.

### Generation of ADC17-70ntΔ strain by CRISPR/CAS9

Guide RNA designed to cut close to the 70-nucleotide region upstream of the ADC17 start codon (agtaacataatgtgctcagcagg) was inserted into pML104 (Addgene number: 67638) to generate pML104-ADC17-70nt. Repair templates to remove the 70-nucleotide upstream region consisting of the flanking region (40 bp) of the 70-nuceotide sequence upstream of ADC17 start codon (f: tcaccaggaaaacaatacttcagaagcttatttctcttgaatgtgctcagcagccggtatcagaagaccaatccagatcg and r: cgatctggattggtcttctgataccggctgctgagcacattcaagagaaataagcttctgaagtattgttttcctggtga) were generated. Additional mutations were inserted to disrupt the PAM site and prevent any further cleavage by Cas9. The repair template was made by annealing (f) and (r) oligo nucleotides in annealing buffer (10 mM (f) oligo, 10 mM (r) oligo, 50 mM NaCl, 10 mM Tris-HCl 7.8, 10 mM MgCl2, and 100 mg/mL BSA). The annealing reaction was incubated at 95°C for 6 min, followed by a gradual decrease of the temperature to 25 °C (1.5 °C/min). pML104-ADC17-70nt was co-transformed in yeast with 10 pmol of annealed repair template, as described above. The clones were verified by PCR using flanking primers and confirmed by sequencing.

### Yeast protein extraction

Yeast samples were pelleted (3200g, 4°C, 4 min). Pellets were washed in 500 μl ice-cold MilliQ water (6200g, 30 sec) and either flash-frozen in dry ice and stored at −20°C for extraction the following day, or extracted immediately. Pellets were resuspended on ice in 400 μl ice-cold 2M LiAc for 1 min, spun down, the supernatant was removed, and pellets resuspended on ice in 400 μl ice-cold 0.4M NaOH for 1 min and spun down again. Pellets were resuspended in 120 μl lysis buffer (0.1 M NaOH, 0.05M EDTA, 2% SDS, 2% beta-mercaptoethanol, PhosStop (Roche), cOmplete protease inhibitor cocktail (Roche)), and boiled for 10 min. 3 μl 4 M acetic acid was added, samples vortexed and boiled for 10 min. Samples were vortexed and spun down (17,000g, 15 min). 80 μl supernatant was added to 20 μl 5x loading buffer (0.25M Tris-HCl pH6.8, 50% Glycerol, 0.05% Bromophenol Blue) and the remainder used to quantify protein concentration using a nanodrop (A280nm, Thermo). Samples were adjusted to the same concentration and stored at −20°C.

### Western blot analyses

Samples were run on homemade 6-14% Bis-Tris acrylamide gels. 2 tubes of 6 ml mix were made up of 6% and 14% acrylamide respectively (0.33M pH 6.5 Bis-Tris, 0.083% APS, 0.083% TEMED). 600 μl of the 14% solution was added to 2x 1 mm Mini-Protean casting gels (biorad). The 14% solution was diluted with 600 μl of the 6% solution and a further 600 μl added to the casting gels. This was repeated until the gels were complete and a comb added. After polymerisation, extracts were loaded to a total of 25-50 μg protein per lane and run at 120 V for 2.5 h at 4°C. Gels were then transferred onto 0.2 μm nitrocellulose membrane (BioRad; 1620112) using a TransBlot Turbo (BioRad) (30 min, 2.5 Amp). Membranes were stained with Ponceau S solution (Santa Cruz, sc-301558), imaged, cut, washed in TBS and blotted for at least 1 h with TBS containing 5% milk, washed 3x in TBS-Tween and incubated with primary antibody overnight. Primary antibody was removed, membranes washed 3x in TBS-T, incubated with secondary antibody for 1 h, washed 3x in TBS-T Tween and imaged on a Chemidoc Touch imaging system (BioRad) using Clarity ECL (BioRad, 170-5061). Where indicated, blots were quantified by densitometry using FIJI with expression of the protein of interest normalised to Ponceau staining (loading control).

### Antibody dilutions

Anti-Adc17 (Bertolotti laboratory; Rabbit; 1:1000), anti-Adc17-(2) (DSTT; Sheep; 1:250; DU66321), anti-Nas6 (Abcam; Rabbit; 1:2000; ab91447), anti-Flag (Sigma Aldrich; Mouse; 1:2000; F3165), anti-Rpt5 (Enzo life sciences; Rabbit; 1:5000; PW8245), anti-20S (Enzo life sciences; Rabbit; 1:2000; PW9355), anti-Mpk1 (Santa Cruz; Mouse; 1:500; sc-374434), anti-p-Mpk1 (Cell Signalling Technology; Rabbit; 1:1000; 9101) and anti-p-Rps6 (Cell Signalling Technology; Rabbit; 1:1000; 2211). Rabbit anti-Adc17 antibody from Bertolotti laboratory was used in all figures excepted for Fig. 1f, 7f, 7j and Extended Data Fig. 4b where Sheep anti-Adc17 antibody was used instead due to stock availability. Anti-mouse IgG, HRP-linked Antibody (Cell Signalling Technology; 1:10000; #7076) and Anti-rabbit IgG, HRP-linked Antibody (Cell Signalling Technology; 1:10000; #7074).

### Proteasome activity assays

Yeasts were grown in YEPD medium overnight at 30°C, then diluted to OD_600nm_ 0.2 in 30 ml YEPD, grown at 32°C to OD_600nm_ 0.5-0.7 and then diluted back to OD_600nm_ 0.2. Treatments were then performed (30 ml with 200 nM rapamycin, 20 ml with 25 μM Latrunculin-B, 20 ml UT control) and cells returned to 32°C for 3 h. 15 OD_600nm_ of cells were spun down (3200g, 4°C, 4 min), resuspended in 800 μl ice-cold water, transferred to a 2 ml tube and spun down again (6,200g, 30 sec, 4°C). The pellet was resuspended in 300 μl Native lysis buffer (50 mM Tris pH 8, 5 mM MgCl2, 0.5 mM EDTA, 5% Glycerol, 1mM DTT, 5 mM ATP) and lysed with 250 μl acid-washed beads (sigma, G-8772) (FastPrep 24, MP, 3 x 30s on, 5 min off). Beads and cell debris were removed by centrifugation (17,000g, 2 min, 4°C), the supernatant was transferred to a fresh tube and centrifuged again (17,000g, 10 min, 4°C). Protein concentration was determined on a nanodrop (A280nm, Thermo) and standardised between samples. 75 μg protein in 1x Native sample buffer (50 mM Tris-HCl pH6.8, 10% glycerol, 0.01% Bromophenol blue) was loaded onto 1.5 mm 3.8-5% acrylamide gradient Native gels (prepared in duplicate as for the western blot gels (above), using 10 ml solutions of 90 mM Tris, 90 mM Boric Acid, 2 mM MgCl2, 1 mM DTT, 0.12% APS, 0.12% TEMED with Acrylamide added to either 3.8% or 5%). Gels were run 2.5 h, 110 V, 4°C in ice-cold native running buffer (0.9 M Tris, 0.9 M Boric Acid, 2 mM MgCl2, 1mM ATP, 1mM DTT). The gels were incubated in 15 ml assay buffer (50 mM Tris-HCl pH 7.5, 150 mM NaCl, 5 mM MgCl2, 10% glycerol) containing 100 μl 10 mM suc-LLVY-AMC fluorogenic substrate (Cambridge biosciences, 4011369), 30°C in the dark, 15-20 min and imaged using a Chemidoc Touch imaging system (BioRad). To image CP assembly, SDS was added to the assay buffer at a final concentration of 0.05% and the gel was re-incubated for 15 minutes prior to imaging again. Gels were transferred onto 0.2 μm nitrocellulose membrane (BioRad; 1620112) using a TransBlot Turbo (BioRad) for western blot analysis.

### Fluorescence microscopy

Yeasts were grown on YEPD plates overnight at 30°C, resuspended to OD_600nm_ 0.2 in YEPD medium and grown at 30°C to OD_600nm_ 0.5-0.7. Cultures were split into 4 ml samples and rapamycin (200 nM final) or Latrunculin-B (25 μM final) were added and samples were returned to 30°C for the indicated time. Formaldehyde (Sigma Aldrich; F8775) was then added to 3.7% and the sample returned to 30°C for 20 min. Samples were spun down (3,200g, 4 min), washed 2x with 7 ml PBS, transferred to a 1.5 ml tube, spun down (7,800g, 2 min), resuspended in 100 μl PBS/0.1% Triton X-100 containing 1:1000 dilution of Rhodamine Phalloidin (Abcam; ab255138) or Phalloidin-iFluor 647 (abcam; ab176759) and incubated at 4°C on a Stuart SB3 vertical rotator in the dark. After 1 h, samples were spun down as before, washed in 1 ml PBS and resuspended in 10 μl ProLong Glass antifade (ThermoFisher; P36980). 3-4 μl were mounted on a SuperFrost microscope slide (VWR; 631-0847), covered with a glass cover slip (VWR; 631-0119) and cured in the dark overnight before imaging on a Zeiss 880 Airyscan microscope (Airyscan mode, Alpha Plan-APO 100x/1.46 Oil DIC VIS objective (Zeiss) and Alpha Plan-APO 63x/1.4 Oil objective (Zeiss)). For the quantification of mRNAs per cell, *adc17Δ* cells with either Ede1 WT or Ede1-aGFP expressing Adc17 mRNA containing MS2 and PP7 stem loops, PCP-GFP and MCP-mCherry were grown and treated with rapamycin as described then fixed and mounted on slides as described, but without phalloidin staining.

For live-cell imaging, 2-3 ml of logarithmically growing yeast cells in DOA media were added to a 35 mm FluoroDish (Fisher Scientific; 15199112) which had been pre-incubated at 30°C with Concanavalin A (Sigma Aldrich, C2010) and allowed to attach for 0.5-1 h. Plates were washed 2x with 2 ml medium to remove unadhered cells and imaged on a Zeiss 880 Airyscan microscope (Airyscan mode, Alpha Plan-APO 100x/1.46 Oil DIC VIS objective (Zeiss, 420792-9800-720)) at 30°C.

All microscopy analysis was carried out using FIJI. For quantification of colocalization of Ede1, Sla1 and Vrp1 with Adc17 mRNA, protein (red) and mRNA (green) punctae (circular punctae, >0.2 μm diameter, >50% brighter than the local cell background) were detected and assessed for colocalization using the ComDet v.0.5.1 plugin using the standard settings. All detected particles were manually checked. For quantification of colocalization of red (translating mRNA) and green (all mRNA) puncta in Suntag experiments, the ComDet v.0.5.1. plugin was used as above. Adc17 mRNA interaction with actin in fixed cells was analysed by performing a maximum intensity Z-projection, then the number of PP7-GFP labelled Adc17 mRNAs (circular punctae, >0.2 μm diameter, >50% brighter than the local cell background) in contact with cortical actin patches (circular punctae, >0.5 μm diameter, 2-fold brighter than actin cables), actin cables (linear structures, 2-fold brighter than cell background), and no actin (remaining Adc17 puncta) was counted manually. To quantify polarity, maximum intensity projections were again performed and budding cells with >6 cortical actin patches in the larger mother cell were counted as depolarised, while those with 6 or under were counted as polarised. To quantify mRNAs per cell, a standard deviation Z-projection was performed and ComDet v.0.5.1 used to detect mRNAs (MCP-mCherry), while cells were counted manually.

### RiboTag RNA isolation

Rpl10-GFP yeasts expressing either FGH17 or FGH17-70ntΔ were grown in YEPD medium overnight at 30°C, then diluted to OD_600nm_ 0.5 in 50 ml YEPD medium and grown at 30°C to OD_600nm_ ~1 and diluted back to OD_600nm_ 0.5 before being treated ± 200nM Rapamycin for 1.5 h at 30°C. After treatment, polysome were stabilised by washing cells in 20 ml ice-cold water containing 0.1 mg/ml CHX before being resuspended in 1 ml RiboTag Lysis Buffer (50 mM Tris, pH 7.5, 100 mM KCl, 12 mM MgCl2, 1% Nonidet P-40, 1 mM DTT, 100U/mL Promega RNasin, 100 mg/mL cycloheximide, cOmplete EDTA-free protease inhibitor cocktail (Roche)) and bead-lysis performed using 500 μl acid-washed beads (Sigma Aldrich, G8772) (FastPrep 24, MP, 5 x 30s on, 5 min off). Ribosome-RNA-containing supernatants were cleared of cell debris by centrifugation (12,000g, 10 min, 4°C). 100 μl slurry GFP binder Sepharose beads (MRC-PPU Reagents) per sample were pre-washed twice in wash buffer (10 mM Tris-HCl, pH7.5, 0.15 M NaCl, 0.5 mM EDTA), before being added to samples and incubated overnight under gentle inversion at 4°C. Beads were washed three times for 10 min with gentle rotation in high-salt buffer (50 mM Tris, pH 7.5, 300 mM KCl, 12 mM MgCl2, 1% Nonidet P-40, 1 mM DTT, 100U/mL Promega RNasin, 100 mg/mL cycloheximide, cOmplete EDTA-free protease inhibitor cocktail (Roche)). RNA was eluted from beads using Qiagen RLT buffer containing 2-Mercaptoethanol and by vortexing 30s. RNA was isolated using RNeasy Kit (74004, QIAGEN) following manufacturer’s instructions, before being analysed by Quantitative RT-PCR.

### Quantitative RT-PCR

Total yeast RNA was extracted using RNeasy Kit (74004, QIAGEN) following manufacturer’s instructions. 1 μg of RNA from untreated and 1.5 h rapamycin treated cells prepared as described above was synthesized into cDNA using SuperScript™ III Reverse Transcriptase (18080093, ThermoFisher). cDNA was diluted 1:10 before the quantitative RT-PCR was performed. Quantitative RT-PCR with primers alg9 (f): cacggatagtggctttggtgaacaattac, alg9 (r): tatgattatctggcagcaggaaagaacttggg, FGH17 (f): gtcctgctggagttcgtgac, FGH17 (r): cgtaatctggaacatcgtatggg, ADC17 (f): cgacgacttggagaacattg, (r): caatgcgtccactctctcat was performed using PowerUp− SYBR− Green Master Mix (A25741; ThermoFisher) on a CFX384 Real-Time PCR Detection system (Biorad). Expression of each gene was normalized to the housekeeping gene ALG9 and expressed as fold change after 1.5h rapamycin treatment calculated using delta-delta Ct method.

### Immunoprecipitation of nascent RPACs using FGH construct

Yeasts expressing FGH or FGH17-70ntΔ were grown in SC-URA medium overnight at 30°C, then diluted to OD_600nm_ 0.2 in 100 ml SC-URA medium and grown at 30°C to OD_600nm_ ~1 and diluted back to OD_600nm_ 0.5 before being treated with 200nM Rapamycin for 1.5 h at 30°C. After treatment, ribosomes were locked on mRNAs by adding 0.1 mg/ml cycloheximide (CHX, final concentration) to the cultures and immediately incubated 10 min on ice, harvested by centrifugation at 4°C (3,200g, 4 min), washed in 20 ml ice-cold water containing 0.1 mg/ml CHX and resuspended in 20 ml of the same solution. Proteins were crosslinked to RNA by treating cells with 254 nm UV (1200 mJ/cm^2^ total; 2x 6000 mJ/cm^2^ with 2 min off in between on ice), then span down as before. Cells were resuspended in 1 ml lysis buffer (0.1 M Tris-HCl pH7.5, 0.5 M LiCl, 10 mM EDTA, 1% Triton-X100, 5 mM DTT, 100 U/ml RNasin (Promega, N2611), cOmplete EDTA-free protease inhibitor cocktail (Roche)) and bead-lysis performed using 500 μl acid-washed beads (Sigma Aldrich, G8772) (4°C, 5x 2 min on, 2 min off using a Disruptor Genie). The supernatant was cleared of cell debris by centrifugation (17,000g, 15 min, 4°C). Protein concentration was determined on a nanodrop (A280nm, Thermo) and protein concentration was standardised between samples. 60 μl M2 anti-flag beads (Sigma Aldrich, M8823) per sample were pre-washed twice in wash buffer (10 mM Tris-HCl, pH7.5, 0.6 M LiCl, 1 mM EDTA), then incubated with 1 mg of lysates for 1 h at 4 °C under rotation. M2-anti-Flag beads were then washed once with lysis buffer, twice with wash buffer. RNase elution was then performed with 100 μl elution buffer (10 mM Tris-HCl pH7.5, 1 mM MgCl2, 40 mM NaCl) containing 5 μl of RNase A/T1 mix (ThermoFisher; EN0551) and placed at 37°C for 1 h under agitation. Elution fractions were subjected to tryptic digestion and tandem mass tag (TMT)-based quantitative proteomics (see below).

### Tryptic digestion of RNase elution

25 μL of protein denaturation buffer (8 M Urea, 50 mM Ammonium bicarbonate pH 8.0 and 5 mM DTT) was added to each RNAse eluent which were denatured at 45 °C for 30 min with gentle shaking (Eppendorf, Thermomixer C, 800 rpm). Samples were centrifuged at 5,000 g for 1 min and cooled to room temperature. Each sample was then incubated with IAA (10 mM final concentration) in the dark at room temperature. Unreacted IAA was then quenched with DTT (5 mM final concentration). Each sample was digested using 0.4 μg trypsin at 37°C overnight and under agitation. The digestion was stopped by adding TFA to the final 0.2% TFA concentration (v/v), centrifuged at 10,000 g for 2 min at room temperature. The supernatant was desalted on ultra microspin column silica C18 (The Nest Group). Desalted peptides were dried using a SpeedVac vacuum centrifuge concentrator (Thermo Fisher) prior to TMT labelling.

### Tandem mass tag (TMT) labelling

Each vacuum-dried sample was resuspended in 50 μL of 100 mM TEAB buffer. The TMT labelling reagents were equilibrated to room temperature and 41 μL anhydrous acetonitrile was added to each reagent channel and gently vortexed for 10 min. 4 μL of each TMT reagent were added to the corresponding sample and labelling performed at room temperature for 1 h with shaking before quenching with 1 μL of 5% hydroxylamine. 2 μL of labelled sample from each channel were analysed by LC-MS/MS to ensure complete labelling prior to mixing. After evaluation, the complete TMT labelled samples were combined, acidified, and dried. The mixture was then desalted with ultra microspin column silica C18 and the eluent from C18 column was dried.

### LC-MS/MS analysis

LC separations were performed with a Thermo Dionex Ultimate 3000 RSLC Nano liquid chromatography instrument. The dried peptides were dissolved in 0.1% formic acid and then loaded on C18 trap column with 3 % ACN/0.1%TFA at a flow rate of 5 μL/min. Peptide separations were performed using EASY-Spray columns (C18, 2 μm, 75 μm × 50 cm) with an integrated nano electrospray emitter at a flow rate of 300 nL/min. Peptides were separated with a 180 min segmented gradient as follows: the first 10 fractions starting from 7%~32% buffer B in 130 min, 32%~45% in 20 min, 45%~95% in 10 min. Peptides eluted from the column were analysed on an Orbitrap Fusion Lumos (Thermo Fisher Scientific, San Jose, CA) mass spectrometer. Spray voltage was set to 2 kV, RF lens level was set at 30%, and ion transfer tube temperature was set to 275°C. The Orbitrap Fusion Lumos was operated in positive ion data dependent mode with high resolution MS2 for reporter ion quantitation. The mass spectrometer was operated in data-dependent Top speed mode with 3 seconds per cycle. The full scan was performed in the range of 350–1500 m/z at nominal resolution of 120 000 at 200 m/z and AGC set to 4 × 10^5^ with maximal injection time 50 ms, followed by selection of the most intense ions above an intensity threshold of 5 × 10^4^ for high collision-induced dissociation (HCD)-MS2 fragmentation in the HCD cell with 38% normalized collision energy. The isolation width was set to 1.2 m/z with no offset. Dynamic exclusion was set to 60 seconds. Monoisotopic precursor selection was set to peptide. Charge states between 2 to 7 were included for MS2 fragmentation. The MS2 scan was performed in the orbitrap using 50 000 resolving power with auto normal range scan from m/z 100 to 500 and AGC target of 5 × 10^4^. The maximal injection time for MS2 scan was set to 120 ms.

### Proteomic data Analysis

All the acquired LC-MS data were analysed using Proteome Discoverer v.2.2 (Thermo Fisher Scientific) with Mascot search engine. Maximum missed cleavage for trypsin digestion was set to 2. Precursor mass tolerance was set to 10 ppm. Fragment ion tolerance was set to 0.02 Da. Carbamidomethylation on cysteine (+57.021 Da) and TMT-10plex tags on N termini as well as lysine (+229.163 Da) were set as static modifications. Variable modifications were set as oxidation on methionine (+15.995 Da) and phosphorylation on serine, threonine, and tyrosine (+79.966 Da). Data were searched against a complete UniProt *S. cerevisiae* (Reviewed 6, 721 entry downloaded Feb 2018). Peptide spectral match (PSM) error rates with a 1% FDR were determined using the forward-decoy strategy modelling true and false matches.

Both unique and razor peptides were used for quantitation. Reporter ion abundances were corrected for isotopic impurities based on the manufacturer’s data sheets. Reporter ions were quantified from MS2 scans using an integration tolerance of 20 ppm with the most confident centroid setting. Signal-to-noise (S/N) values were used to represent the reporter ion abundance with a co-isolation threshold of 50% and an average reporter S/N threshold of 10 and above required for quantitation from each MS2 spectra to be used. The S/N value of each reporter ions from each PSM were used to represent the abundance of the identified peptides. The summed abundance of quantified peptides was used for protein quantitation. The total peptide amount was used for the normalisation. Protein ratios were calculated from medians of summed sample abundances of replicate groups. Standard deviation was calculated from all biological replicate values. The standard deviation of all biological replicates lower than 25% were used for further analyses. Multiple unpaired t-test was used to determine the significant differences between FGH or FGH17-70ntΔ.

### Statistical analysis

Each experiment was repeated independently a minimum of three times, as indicated. The standard deviation (s.d.) of the mean of at least four independent experiments is shown in the graphs, or as indicated. ns (not significant) or asterisks *(*P* ≤ 0.05; \*\**P* ≤ 0.01; \*\*\**P* ≤ 0.001) indicate *P* values obtained from one-way analysis of variance (ANOVA) (Dunnett multiple comparison test: Fig. 5f, 5h and 6b), two-way ANOVA (Tukey multiple comparison test: Fig. 3f, 4c, 7c, 7g, 7i and 7k, and Ext. Data Fig. 5e) or unpaired t-test (Fig. 1b, 1e and 3c, and Ext. Data Fig. 1c, 3a and 3b) to probe for statistical significance. No statistical method was used to predetermine sample size.

### Data availability

All the data generated or analysed during the current study are included in this published article and its supplementary files (Supplementary information and Source data). The datasets generated during and/or analysed during the current study are available from the corresponding author on reasonable request. The mass spectrometry proteomics data have been deposited to the ProteomeXchange Consortium via the PRIDE partner repository with the dataset identifier PXD027655.

## References

1. Hipp, M. S., Kasturi, P. & Hartl, F. U. The proteostasis network and its decline in ageing. Nat. Rev. Mol. Cell Biol. 20, 421–435 (2019).

2. Labbadia, J. & Morimoto, R. I. The Biology of Proteostasis in Aging and Disease. Annu. Rev. Biochem. 84, 435–464 (2015).

3. Pilla, E., Schneider, K. & Bertolotti, A. Coping with Protein Quality Control Failure. Annu. Rev. Cell Dev. Biol. 33, 439–465 (2017).

4. Rousseau, A. & Bertolotti, A. Regulation of proteasome assembly and activity in health and disease. Nat. Rev. Mol. Cell Biol. 19, 697–712 (2018).

5. Dikic, I. Proteasomal and Autophagic Degradation Systems. Annu. Rev. Biochem. 86, 193–224 (2017).

6. Yu, H. & Matouschek, A. Recognition of Client Proteins by the Proteasome. Annu. Rev. Biophys. 46, 149–173 (2017).

7. Budenholzer, L., Cheng, C. L., Li, Y. & Hochstrasser, M. Proteasome Structure and Assembly. J. Mol. Biol. 429, 3500–3524 (2017).

8. Ben-Sahra, I. & Manning, B. D. mTORC1 signaling and the metabolic control of cell growth. Curr. Opin. Cell Biol. 45, 72–82 (2017).

9. González, A. & Hall, M. N. Nutrient sensing and TOR signaling in yeast and mammals. EMBO J. 36, 397–408 (2017).

10. Saxton, R. A. & Sabatini, D. M. mTOR Signaling in Growth, Metabolism, and Disease. Cell 168, 960–976 (2017).

11. Noda, T. Regulation of Autophagy through TORC1 and mTORC1. Biomolecules 7, (2017).

12. Zhao, J., Zhai, B., Gygi, S. P. & Goldberg, A. L. mTOR inhibition activates overall protein degradation by the ubiquitin proteasome system as well as by autophagy. Proc. Natl. Acad. Sci. 112, 15790–15797 (2015).

13. Rousseau, A. & Bertolotti, A. An evolutionarily conserved pathway controls proteasome homeostasis. Nature 536, 184–189 (2016).

14. Suraweera, A., Münch, C., Hanssum, A. & Bertolotti, A. Failure of Amino Acid Homeostasis Causes Cell Death following Proteasome Inhibition. Mol. Cell 48, 242–253 (2012).

15. Vabulas, R. M. & Hartl, F. U. Protein Synthesis upon Acute Nutrient Restriction Relies on Proteasome Function. Science 310, 1960–1963 (2005).

16. Torres, J., Di Como, C. J., Herrero, E. & de la Torre-Ruiz, M. A. Regulation of the Cell Integrity Pathway by Rapamycin-sensitive TOR Function in Budding Yeast. J. Biol. Chem. 277, 43495–43504 (2002).

17. Waite, K. A., Burris, A., Vontz, G., Lang, A. & Roelofs, J. Proteasome autophagy is specifically regulated and requires factors dispensible for general autophagy. bioRxiv 2021.03.26.437055 (2021) doi:10.1101/2021.03.26.437055.

18. Sanz, E. et al. Cell-type-specific isolation of ribosome-associated mRNA from complex tissues. Proc. Natl. Acad. Sci. 106, 13939–13944 (2009).

19. Funakoshi, M., Tomko Jr., R. J., Kobayashi, H. & Hochstrasser, M. Multiple Assembly Chaperones Govern Biogenesis of the Proteasome Regulatory Particle Base. Cell 137, 887–899 (2009).

20. Kaneko, T. et al. Assembly Pathway of the Mammalian Proteasome Base Subcomplex Is Mediated by Multiple Specific Chaperones. Cell 137, 914–925 (2009).

21. Roelofs, J. et al. Chaperone-mediated pathway of proteasome regulatory particle assembly. Nature 459, 861–865 (2009).

22. Shchepachev, V. et al. Defining the RNA interactome by total RNA-associated protein purification. Mol. Syst. Biol. 15, e8689 (2019).

23. Wang, C., Han, B., Zhou, R. & Zhuang, X. Real-time imaging of translation on single mRNA transcripts in live cells. Cell 165, 990–1001 (2016).

24. Toshima, J. Y. et al. Spatial dynamics of receptor-mediated endocytic trafficking in budding yeast revealed by using fluorescent α-factor derivatives. Proc. Natl. Acad. Sci. 103, 5793–5798 (2006).

25. Munn, A. L. & Thanabalu, T. Verprolin: a cool set of actin-binding sites and some very HOT prolines. IUBMB Life 61, 707–712 (2009).

26. Goode, B. L., Eskin, J. A. & Wendland, B. Actin and Endocytosis in Budding Yeast. Genetics 199, 315–358 (2015).

27. Lu, R., Drubin, D. G. & Sun, Y. Clathrin-mediated endocytosis in budding yeast at a glance. J. Cell Sci. 129, 1531–1536 (2016).

28. Song, J. et al. Essential Genetic Interactors of SIR2 Required for Spatial Sequestration and Asymmetrical Inheritance of Protein Aggregates. PLOS Genet. 10, e1004539 (2014).

29. Mulet, J. M., Martin, D. E., Loewith, R. & Hall, M. N. Mutual Antagonism of Target of Rapamycin and Calcineurin Signaling. J. Biol. Chem. 281, 33000–33007 (2006).

30. Irazoqui, J. E., Howell, A. S., Theesfeld, C. L. & Lew, D. J. Opposing Roles for Actin in Cdc42p Polarization. Mol. Biol. Cell 16, 1296–1304 (2005).

31. Su, K.-H. & Dai, C. mTORC1 senses stresses: Coupling stress to proteostasis. BioEssays News Rev. Mol. Cell. Dev. Biol. 39, (2017).

32. Advani, V. M. & Ivanov, P. Translational control under stress: reshaping the translatome. BioEssays News Rev. Mol. Cell. Dev. Biol. 41, e1900009 (2019).

33. Pan, T. Adaptive Translation as a Mechanism of Stress Response and Adaptation. Annu. Rev. Genet. 47, 121–137 (2013).

34. Yamasaki, S. & Anderson, P. Reprogramming mRNA translation during stress. Curr. Opin. Cell Biol. 20, 222–226 (2008).

35. Medioni, C., Mowry, K. & Besse, F. Principles and roles of mRNA localization in animal development. Dev. Camb. Engl. 139, 3263–3276 (2012).

36. Herbert, S. P. & Costa, G. Sending messages in moving cells: mRNA localization and the regulation of cell migration. Essays Biochem. 63, 595–606 (2019).

37. Glock, C., Heumüller, M. & Schuman, E. M. mRNA transport & local translation in neurons. Curr. Opin. Neurobiol. 45, 169–177 (2017).

38. Kraut-Cohen, J. et al. Translation-and SRP-independent mRNA targeting to the endoplasmic reticulum in the yeast Saccharomyces cerevisiae. Mol. Biol. Cell 24, 3069–3084 (2013).

39. Casolari, J. M. et al. Widespread mRNA Association with Cytoskeletal Motor Proteins and Identification and Dynamics of Myosin-Associated mRNAs in S. cerevisiae. PLOS ONE 7, e31912 (2012).

40. Medina-Munoz, H. C., Lapointe, C. P., Porter, D. F. & Wickens, M. Records of RNA locations in living yeast revealed through covalent marks. Proc. Natl. Acad. Sci. 117, 23539–23547 (2020).

41. Takizawa, P. A., Sil, A., Swedlow, J. R., Herskowitz, I. & Vale, R. D. Actin-dependent localization of an RNA encoding a cell-fate determinant in yeast. Nature 389, 90–93 (1997).

42. Reid, D. W. & Nicchitta, C. V. Diversity and selectivity in mRNA translation on the endoplasmic reticulum. Nat. Rev. Mol. Cell Biol. 16, 221–231 (2015).

43. Lesnik, C., Golani-Armon, A. & Arava, Y. Localized translation near the mitochondrial outer membrane: An update. RNA Biol. 12, 801–809 (2015).

44. Rangaraju, V., tom Dieck, S. & Schuman, E. M. Local translation in neuronal compartments: how local is local? EMBO Rep. 18, 693–711 (2017).

45. Vedula, P. et al. Different translation dynamics of β-and γ-actin regulates cell migration. bioRxiv 2021.01.05.425467 (2021) doi:10.1101/2021.01.05.425467.

46. Sundell, C. L. & Singer, R. H. Requirement of microfilaments in sorting of actin messenger RNA. Science 253, 1275–1277 (1991).

47. Katz, Z. B. et al. β-Actin mRNA compartmentalization enhances focal adhesion stability and directs cell migration. Genes Dev. 26, 1885–1890 (2012).

48. Scholz, D. et al. Microtubule-dependent distribution of mRNA in adult cardiocytes. Am. J. Physiol. Heart Circ. Physiol. 294, H1135–1144 (2008).

49. Wilfling, F. et al. A Selective Autophagy Pathway for Phase-Separated Endocytic Protein Deposits. Mol. Cell 80, 764–778. e7 (2020).

50. Fricke, B., Heink, S., Steffen, J., Kloetzel, P.-M. & Krüger, E. The proteasome maturation protein POMP facilitates major steps of 20S proteasome formation at the endoplasmic reticulum. EMBO Rep. 8, 1170–1175 (2007).

51. Svitkina, T. M. Actin Cell Cortex: Structure and Molecular Organization. Trends Cell Biol. 30, 556–565 (2020).

52. van Bommel, B., Konietzny, A., Kobler, O., Bär, J. & Mikhaylova, M. F-actin patches associated with glutamatergic synapses control positioning of dendritic lysosomes. EMBO J. 38, e101183 (2019).

53. Kumari, S. et al. Actin foci facilitate activation of the phospholipase C-γ in primary T lymphocytes via the WASP pathway. eLife 4, e04953 (2015).

54. Basu, S. & Lamprecht, R. The Role of Actin Cytoskeleton in Dendritic Spines in the Maintenance of Long-Term Memory. Front. Mol. Neurosci. 11, (2018).

55. Nakahata, Y. & Yasuda, R. Plasticity of Spine Structure: Local Signaling, Translation and Cytoskeletal Reorganization. Front. Synaptic Neurosci. 10, (2018).

56. Honkura, N., Matsuzaki, M., Noguchi, J., Ellis-Davies, G. C. R. & Kasai, H. The Subspine Organization of Actin Fibers Regulates the Structure and Plasticity of Dendritic Spines. Neuron 57, 719–729 (2008).

57. Hamilton, A. M. et al. Activity-Dependent Growth of New Dendritic Spines Is Regulated by the Proteasome. Neuron 74, 1023–1030 (2012).

58. Bingol, B. & Schuman, E. M. Activity-dependent dynamics and sequestration of proteasomes in dendritic spines. Nature 441, 1144–1148 (2006).

59. Smethurst, D. G. J., Dawes, I. W. & Gourlay, C. W. Actin – a biosensor that determines cell fate in yeasts. FEMS Yeast Res. 14, 89–95 (2014).

60. Janke, C. et al. A versatile toolbox for PCR-based tagging of yeast genes: new fluorescent proteins, more markers and promoter substitution cassettes. Yeast 21, 947–962 (2004).

