## Extended data and supplementary tables for "Actin remodelling controls proteasome homeostasis upon stress"

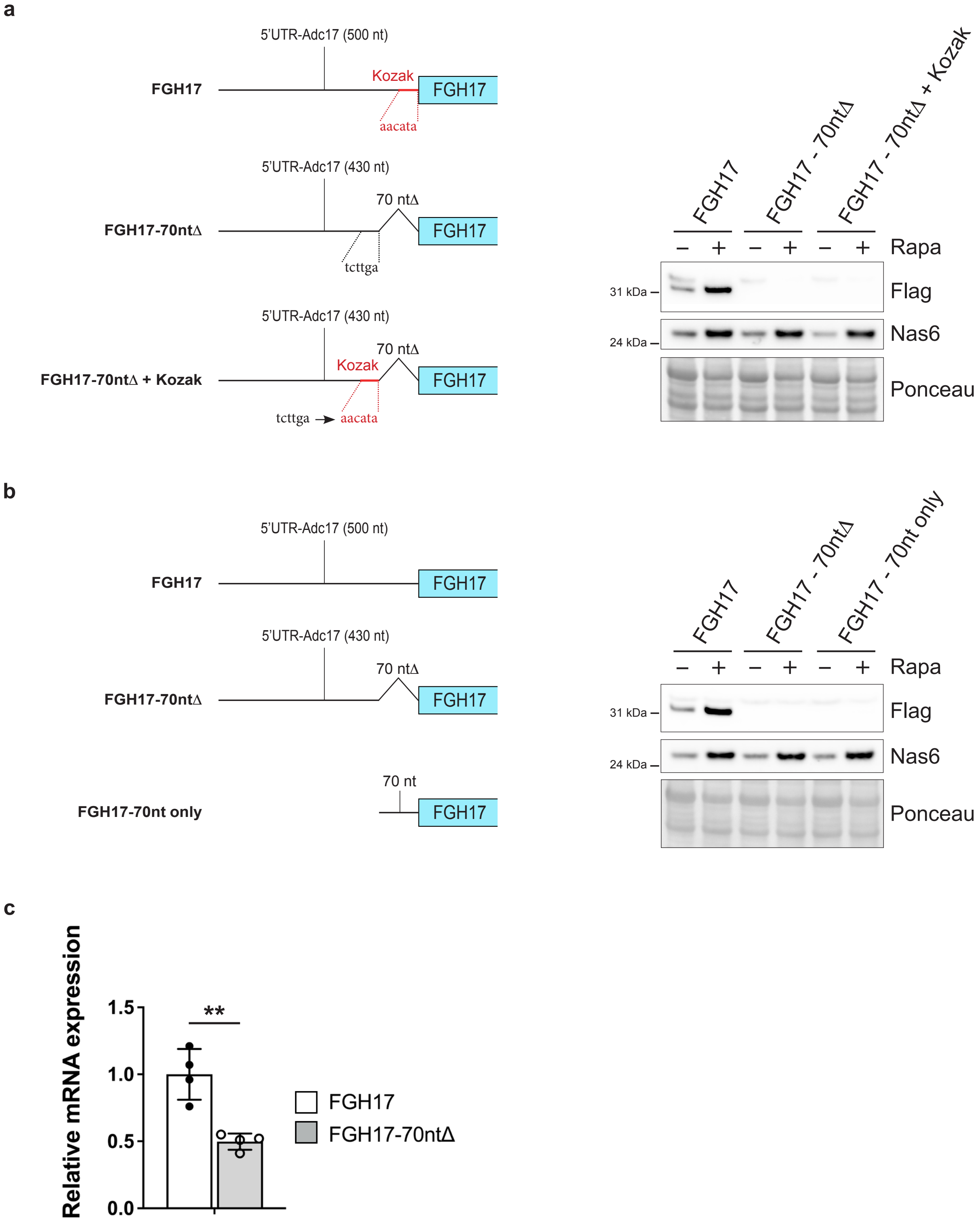


**Extended data Fig. 1| Characterisation of the region important for FGH17 regulation**

**Extended data Fig. 1| Characterisation of the region important for FGH17 regulation. a,** Schematic showing the introduction of the ADC17 5'UTR Kozak sequence into the FGH17-70nt$\Delta$ vector. Western blot analysis showing the impact on FGH17 levels in cells treated ± 200 nM rapamycin (Rapa) for 4 h. Ponceau S staining was used as a loading control. **b,** FGH17 vectors with the full 5'UTR, the 5'UTR lacking the 70 nucleotides upstream of the start codon (FGH17-70nt$\Delta$) or containing only the 70 nucleotides upstream of the start codon (FGH17-70nt only). Western blot analysis showing the impact on FGH17 levels in cells treated ± 200 nM rapamycin (Rapa) for 4 h. Ponceau S staining was used as a loading control. **c,** Relative abundance of FGH17 and FGH17-70nt$\Delta$ mRNA (mRNA of interest normalised to ALG9 housekeeping mRNA) in yeast cells. For each bar, $n$=4 biologically independent experiments for each condition. Statistical analysis was carried out using unpaired t-test. $^{**}P \leq 0.01$. **a, b,** Data are representative of three independent biological replicates.

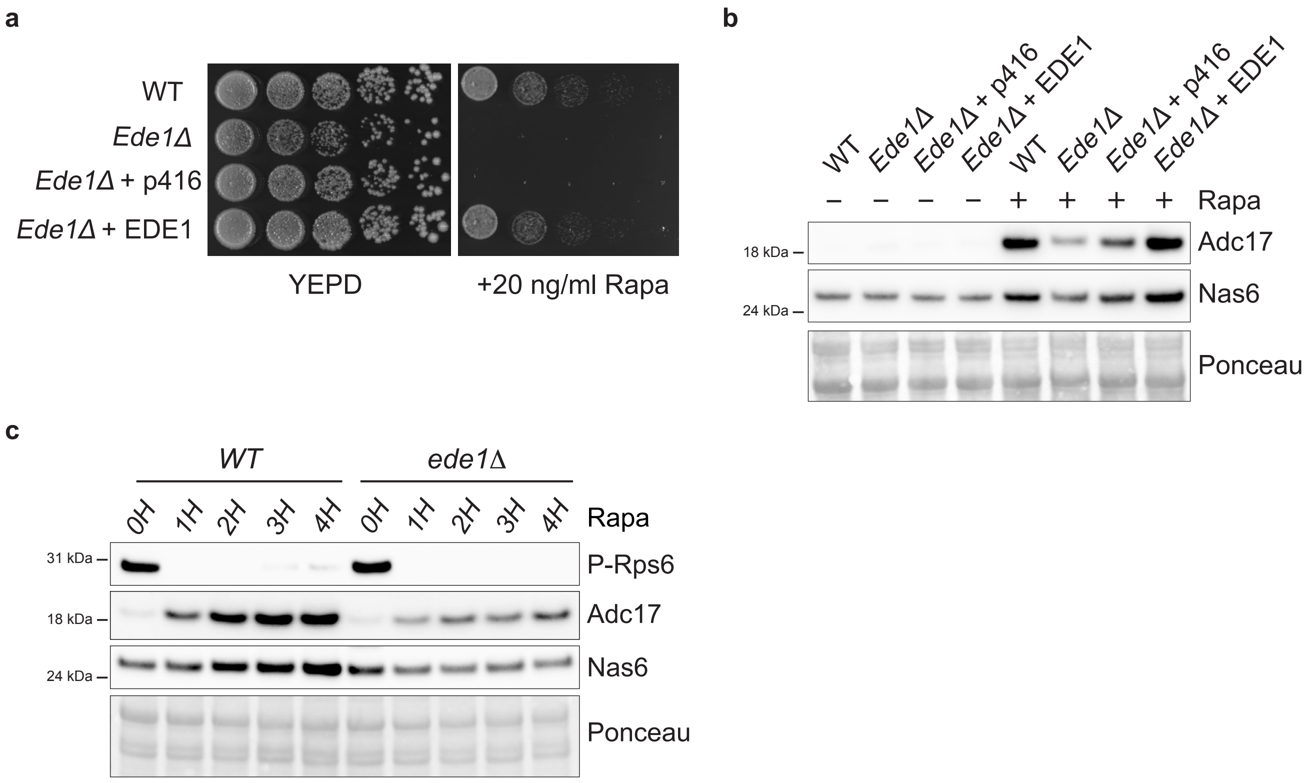


**Extended data Fig. 2| Ede1 regulates RPAC levels downstream of TORC1 inhibition**

**Extended data Fig. 2| Ede1 regulates RPAC levels downstream of TORC1 inhibition. a,** Cells spotted in a fivefold dilution and grown for 3 days on plates ± 20 ng/ml rapamycin. **b**, Western blot analysis of RPACs in WT and *ede1*Δ cells treated ± 200 nM rapamycin (Rapa) for 4 h. Ponceau S staining was used as a loading control. **c**, Western blot analysis of RPACs and P-Rps6 in WT and *ede1*Δ cells treated ± 200 nM rapamycin (Rapa) for the indicated time. Ponceau S staining was used as a loading control. **a-c**, Data are representative of three independent biological replicates.

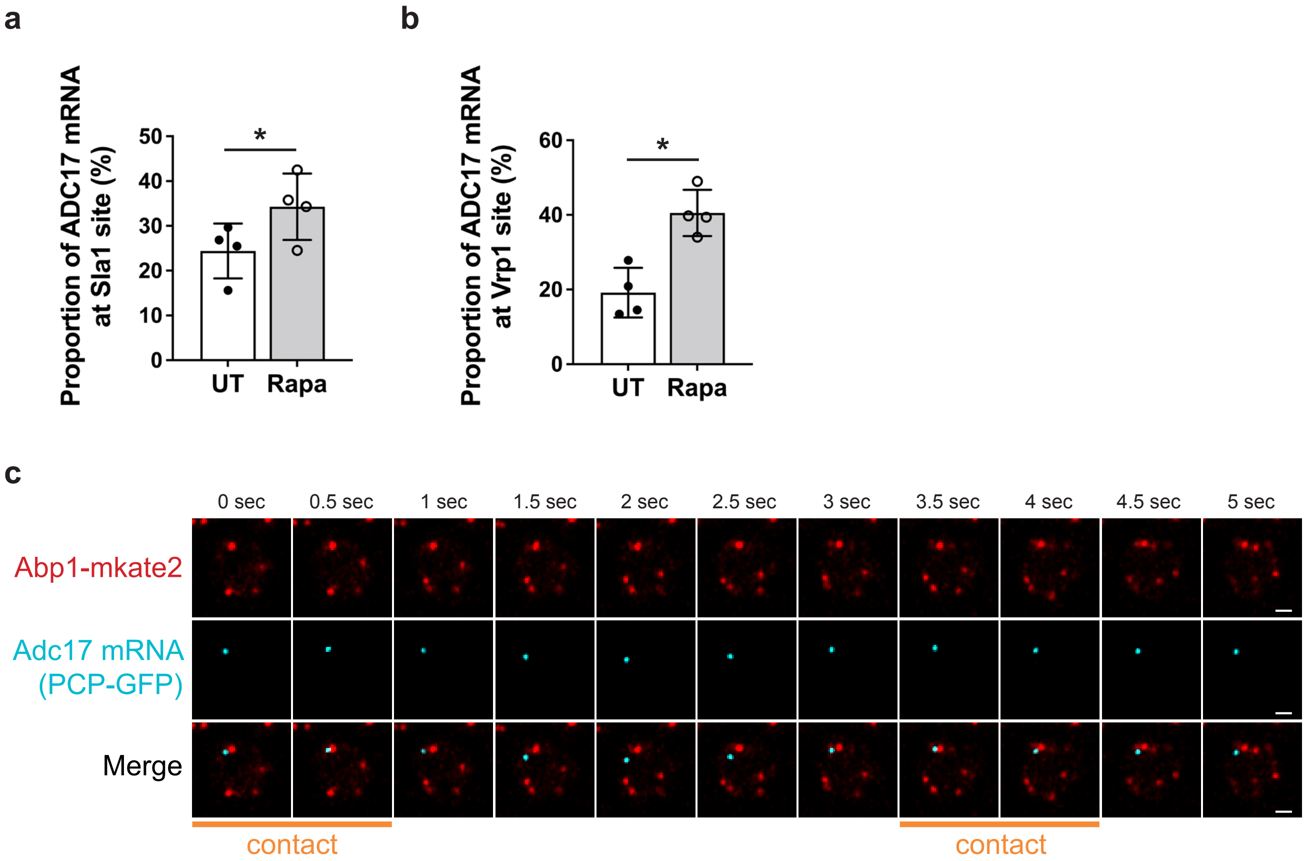


**Extended data Fig. 3| ADC17 mRNA interacts with cortical actin patch proteins**

**Extended data Fig. 3| ADC17 mRNA interacts with cortical actin patch proteins. a,** Frequency of ADC17 mRNAs colocalizing with Sla1-mKate2 in cells grown for 3 h ± 200 nM rapamycin (Rapa). Untreated (UT). For each bar, *n*=4 biologically independent experiments with at least 500 ADC17 mRNAs for each condition. Statistical analysis was carried out using unpaired t-test. $^{**}P \leq 0.01$. **b,** Frequency of ADC17 mRNAs colocalizing with Vrp1-mKate2 in cells grown for 3 h ± 200 nM rapamycin (Rapa). Untreated (UT). For each bar, *n*=4 biologically independent experiments with at least 400 ADC17 mRNAs for each condition. Statistical analysis was carried out using unpaired t-test. $^{**}P \leq 0.01$. **c,** Representative single frames from time-lapse imaging showing contacts between Abp1-mKate2 (red) and PCP-GFP-labelled ADC17 mRNA (cyan). Scale bars, 1 μm.

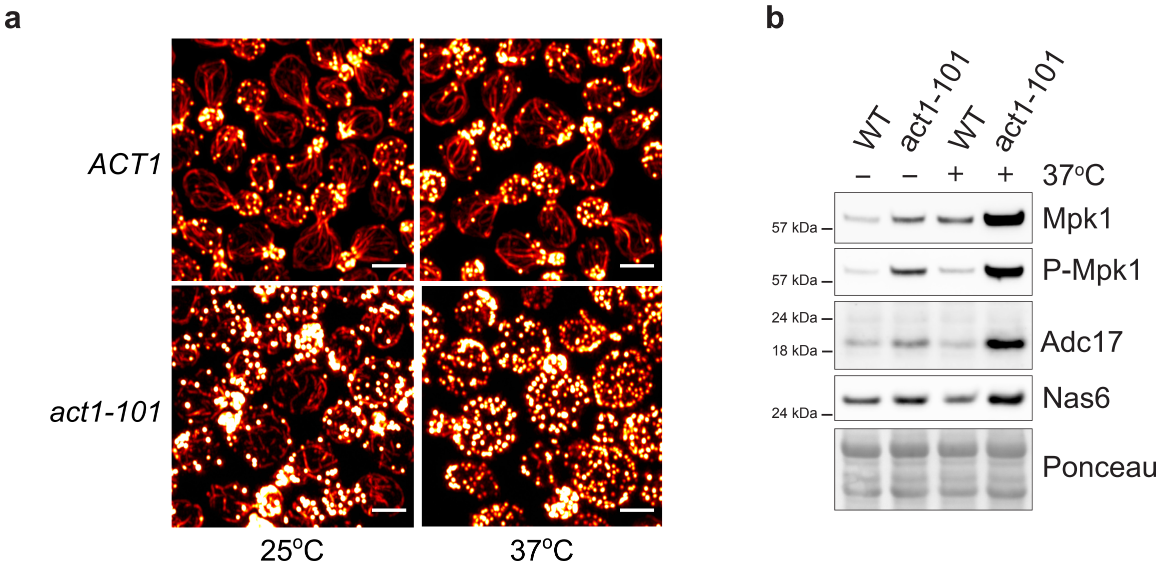


**Extended data Fig. 4| Genetic disruption of actin induces RPAC levels**

**Extended data Fig. 4| Genetic disruption of actin induces RPAC levels. a,** Representative microscopy images (maximum intensity Z-projection) of WT (ACT1) and *act1-101* cells either grown at the permissive temperature (25°C) or shifted to the non-permissive temperature (37°C) for 4 h and stained with Rhodamine phalloidin to visualise actin (hot red LUT). Scale bars, 3 μm. **b,** Western blot analysis of RPACs in WT and *act1-101* cells either grown at the permissive temperature (25°C) or shifted to the non-permissive temperature (37°C) for 4 h. Ponceau S staining was used as a loading control. Data are representative of three independent biological replicates.

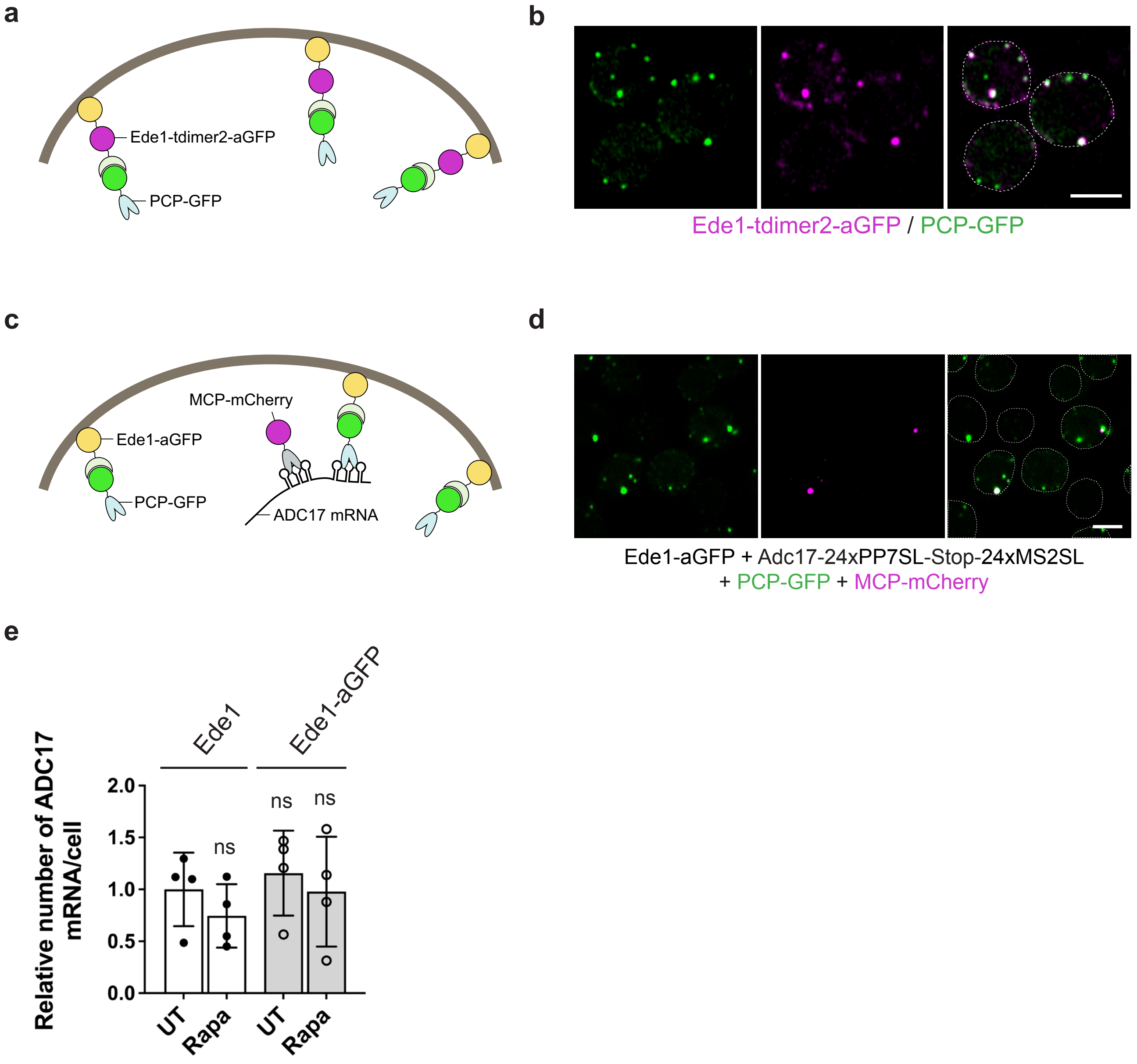


**Extended data Fig. 5| Validation of ADC17 mRNA tethering to Ede1**

**Extended data Fig. 5| Validation of ADC17 mRNA tethering to Ede1. a,** Schematic representation of PCP-GFP recruitment to Ede1 fused with tdimer2 and aGFP (nanobody against GFP). **b**, Representative microscopy images of yeast cells containing PCP-GFP (green) and Ede1-tdimer2 (magenta) tagged with a nanobody against GFP (Ede1-tdimer2-aGFP). Scale bars, 3 μm. **c,** Schematic representation of doubly tagged ADC17 mRNA (magenta) recruitment to Ede1-aGFP/PCP-GFP complex. **d**, Representative microscopy images of yeast cells expressing PCP-GFP (green), MCP-mCherry (magenta), Ede1 tagged with a nanobody against GFP (Ede1-aGFP) and ADC17 mRNA containing PP7-stem-loops (PP7SL) and MS2-stem-loops (MS2SL) which are recognised by PCP-GFP and MCP-mCherry, respectively. Scale bars, 3 μm. **e,** Quantification of the number of ADC17 mRNAs per cell in *adc17*Δ Ede1 WT and Ede1-aGFP cells expressing *ADC17* mRNA containing MS2 and PCP stem loops, PP7-GFP and MCP-mCherry grown for 2 h ± 200 nM rapamycin (Rapa). Untreated (UT). mRNAs were quantified using MCP-mCherry as a marker. For each bar, *n*=4 biologically independent experiments with at least 250 ADC17 mRNAs for each condition. Statistical analysis was carried out using two-way ANOVA t-test (Tukey multiple comparison test). ns, not significant.

**Supplementary video 1,2.** Representative time-lapse recording showing contacts between Ede1-GFPEnvy (green) and ADC17 mRNA (magenta). ADC17 mRNAs are labelled with PP7 bacteriophage coat protein (PCP) fused to mKate2 (magenta). The experiments were repeated at least three times. Scale bars, 3 μm.

**Supplementary video 3,4.** Representative time-lapse recording of yeast cells expressing ADC17-SunTag reporter mRNA. Translating ADC17 mRNAs are GFP (green)- and mCherry (magenta)-positive while non-translating ADC17 mRNAs are only positive for GFP. The experiments were repeated at least three times. Scale bars, 3 μm.

**Supplementary video 5,6.** Representative time-lapse recording showing the transport of ADC17 mRNA along actin cable. Actin cable and ADC17 mRNA are shown in red (Abp140-mKate2) and cyan (PCP-GFP), respectively. The experiments were repeated at least three times. Scale bars, 3 μm.

**Supplementary video 7,8.** Representative time-lapse recording showing contacts between the cortical actin patch marker Abp1-mKate2 (red) and PCP-GFP-labelled ADC17 mRNA (cyan). The experiments were repeated at least three times. Scale bars, 3 μm.

**Supplementary Table 1.** List of the *S. cerevisiae* strains used in this study.

**Supplementary Table 1 | List of strains used in this study.**

| **Strain** | **Genotype**; plasmids in brakcets | **Source** |
|---|---|---|
| BY4741 | *MATa his3Δ1 leu2Δ0 met15Δ0 ura3Δ0* | Horizon Discovery |
| BY4741 + FGH17 | *MATa his3Δ1 leu2Δ0 met15Δ0 ura3Δ0 [p416:FGH17::URA3]* | This study |
| BY4741 + FGH17-5'UTRΔ | *MATa his3Δ1 leu2Δ0 met15Δ0 ura3Δ0 [p416:FGH17-5'UTR Δ::URA3]* | This study |
| BY4741 + FGH17-3'UTRΔ | *MATa his3Δ1 leu2Δ0 met15Δ0 ura3Δ0 [p416:FGH17-3'UTR Δ::URA3]* | This study |
| BY4741 + FGH17-70ntΔ | *MATa his3Δ1 leu2Δ0 met15Δ0 ura3Δ0 [p416:FGH17-70nt Δ::URA3]* | This study |
| BY4741 + FGH17-40ntΔ | *MATa his3Δ1 leu2Δ0 met15Δ0 ura3Δ0 [p416:FGH17-40nt Δ::URA3]* | This study |
| BY4741 + FGH17-23ntΔ | *MATa his3Δ1 leu2Δ0 met15Δ0 ura3Δ0 [p416:FGH17-23nt Δ::URA3]* | This study |
| rps6AΔ | *MATa his3Δ1 met15Δ0 ura3Δ0 rps6a::LEU2* | This study |
| rps18BΔ | *MATa his3Δ1 met15Δ0 ura3Δ0 rps18b::LEU2* | This study |
| ede1Δ | *MATa his3Δ1 leu2Δ0 met15Δ0 ura3Δ0 ede1::kanMx* | Horizon Discovery |
| cup1Δ | *MATa his3Δ1 leu2Δ0 met15Δ0 ura3Δ0 cup1-1/2::kanMx* | Horizon Discovery |
| Adc17-24xPP7SL + PCP-mKate2 | *MATa his3Δ1 leu2Δ0 met15Δ0 ura3Δ0 ADC17-24xPP7SL-LoxP [pFA6:cyc1p-PCP-mKate2::HIS3]* | This study |
| Adc17-24xPP7SL Ede1-3xHA-GFPEnvy + PCP-mKate2 | *MATa his3Δ1 leu2Δ0 met15Δ0 ura3Δ0 ADC17-24xPP7SL-LoxP EDE1-3xHA-GFPENVY:KanMx [pFA6:cyc1p-PCP-mKate2::HIS3]* | This study |
| Adc17-SunTag | *MATa his3Δ1 leu2Δ0 met15Δ0 ura3Δ0 [p416:adc17p-Adc17-SunTag(24x)-PP7SL(24x)::URA3 + pFA6:cyc1p-PCP-EGFP(2X)-cyc1p-scFV-GCN4-mCherry::HIS3 + ]* | This study |
| Adc17-SunTag ede1Δ | *MATa his3Δ1 leu2Δ0 met15Δ0 ura3Δ0 ede1::KanMx [p416:adc17p-Adc17-SunTag(24x)-PP7SL(24x)::URA3 + pFA6:cyc1p-PCP-EGFP(2X)-cyc1p-scFV-GCN4-mCherry::HIS3 +* | This study |
| syp1Δ | *MATa his3Δ1 leu2Δ0 met15Δ0 ura3Δ0 syp1::kanMx* | Horizon Discovery |
| clc1Δ | *MATa his3Δ1 leu2Δ0 met15Δ0 ura3Δ0 clc1::kanMx* | Horizon Discovery |
| chc1Δ | *MATa his3Δ1 leu2Δ0 met15Δ0 ura3Δ0 chc1::kanMx* | Horizon Discovery |
| pal1Δ | *MATa his3Δ1 leu2Δ0 met15Δ0 ura3Δ0 pal1::kanMx* | Horizon Discovery |
| yap1801Δ | *MATa his3Δ1 leu2Δ0 met15Δ0 ura3Δ0 yap1801::kanMx* | Horizon Discovery |
| yap1802Δ | *MATa his3Δ1 leu2Δ0 met15Δ0 ura3Δ0 yap1802::kanMx* | Horizon Discovery |
| alp1Δ | *MATa his3Δ1 leu2Δ0 met15Δ0 ura3Δ0 alp1::kanMx* | Horizon Discovery |
| alp3Δ | *MATa his3Δ1 leu2Δ0 met15Δ0 ura3Δ0 alp3::kanMx* | Horizon Discovery |
| aps2Δ | *MATa his3Δ1 leu2Δ0 met15Δ0 ura3Δ0 aps2::kanMx* | Horizon Discovery |
| apm4Δ | *MATa his3Δ1 leu2Δ0 met15Δ0 ura3Δ0 apm4::kanMx* | Horizon Discovery |
| ent1Δ | *MATa his3Δ1 leu2Δ0 met15Δ0 ura3Δ0 ent1::kanMx* | Horizon Discovery |
| ent2Δ | *MATa his3Δ1 leu2Δ0 met15Δ0 ura3Δ0 ent2::kanMx* | Horizon Discovery |
| end3Δ | *MATa his3Δ1 leu2Δ0 met15Δ0 ura3Δ0 end3::kanMx* | Horizon Discovery |
| sla1Δ | *MATa his3Δ1 leu2Δ0 met15Δ0 ura3Δ0 sla1::kanMx* | Horizon Discovery |
| lsb3Δ | *MATa his3Δ1 leu2Δ0 met15Δ0 ura3Δ0 lsb3::kanMx* | Horizon Discovery |
| lsb4Δ | *MATa his3Δ1 leu2Δ0 met15Δ0 ura3Δ0 lsb4::kanMx* | Horizon Discovery |
| lsb5Δ | *MATa his3Δ1 leu2Δ0 met15Δ0 ura3Δ0 lsb5::kanMx* | Horizon Discovery |
| ubx3Δ | *MATa his3Δ1 leu2Δ0 met15Δ0 ura3Δ0 ubx3::kanMx* | Horizon Discovery |
| gts1Δ | *MATa his3Δ1 leu2Δ0 met15Δ0 ura3Δ0 gst1::kanMx* | Horizon Discovery |
| ldb17Δ | *MATa his3Δ1 leu2Δ0 met15Δ0 ura3Δ0 ldb17::kanMx* | Horizon Discovery |
| bbc1Δ | *MATa his3Δ1 leu2Δ0 met15Δ0 ura3Δ0 bbc1::kanMx* | Horizon Discovery |
| aim21Δ | *MATa his3Δ1 leu2Δ0 met15Δ0 ura3Δ0 aim21::kanMx* | Horizon Discovery |
| ubp7Δ | *MATa his3Δ1 leu2Δ0 met15Δ0 ura3Δ0 ubp7::kanMx* | Horizon Discovery |
| bzz1Δ | *MATa his3Δ1 leu2Δ0 met15Δ0 ura3Δ0 bzz1::kanMx* | Horizon Discovery |
| vrp1Δ | *MATa his3Δ1 leu2Δ0 met15Δ0 ura3Δ0 vrp1::kanMx* | Horizon Discovery |
| myo3Δ | *MATa his3Δ1 leu2Δ0 met15Δ0 ura3Δ0 myo3::kanMx* | Horizon Discovery |
| myo5Δ | *MATa his3Δ1 leu2Δ0 met15Δ0 ura3Δ0 myo5::kanMx* | Horizon Discovery |
| rvs161Δ | *MATa his3Δ1 leu2Δ0 met15Δ0 ura3Δ0 rvs161::kanMx* | Horizon Discovery |
| rvs167Δ | *MATa his3Δ1 leu2Δ0 met15Δ0 ura3Δ0 rvs167::kanMx* | Horizon Discovery |
| vps1Δ | *MATa his3Δ1 leu2Δ0 met15Δ0 ura3Δ0 vps1::kanMx* | Horizon Discovery |
| Adc17-24xPP7SL + PCP-GFP(2x) | *MATa his3Δ1 leu2Δ0 met15Δ0 ura3Δ0 ADC17-24xPP7SL-LoxP [pFA6:cyc1p-PCP-EGFP(2X)::HIS3]* | This study |
| Adc17-24xPP7SL Abp140-3xHA-mKate2 + PCP-EGFP(2X) | *MATa his3Δ1 leu2Δ0 met15Δ0 ura3Δ0 ADC17-24xPP7SL-LoxP ABP140-3xHA-mKATE2:KanMx [pFA6:cyc1p-PCP-EGFP(2X)::HIS3]* | This study |
| Adc17-24xPP7SL Abp1-3xHA-mKate2 + PCP-EGFP(2X) | *MATa his3Δ1 leu2Δ0 met15Δ0 ura3Δ0 ADC17-24xPP7SL-LoxP ABP1-3xHA-mKATE2:KanMx [pFA6:cyc1p-PCP-EGFP(2X)::HIS3]* | This study |
| Adc17-24xPP7SL ede1Δ + PCP-EGFP(2X) | *MATa his3Δ1 leu2Δ0 met15Δ0 ura3Δ0 ADC17-24xPP7SL-LoxP ede1Δ::LEU2 [pFA6:cyc1p-PCP-EGFP(2X)::HIS3]* | This study |
| Adc17-24xPP7SL Ede1-aGFP + PCP-GFP(2x) | *MATa his3Δ1 leu2Δ0 met15Δ0 ura3Δ0 ADC17-24xPP7SL-LoxP EDE1-aGFP:LEU2 [pFA6:cyc1p-PCP-EGFP(2X)::HIS3]* | This study |
| ede1Δ + p416 | *MATa his3Δ1 leu2Δ0 met15Δ0 ura3Δ0 ede1::kanMx [p416]* | This study |
| ede1Δ + p416-Ede1 | *MATa his3Δ1 leu2Δ0 met15Δ0 ura3Δ0 ede1::kanMx [p416-ede1p-Ede1]* | This study |
| Adc17-24xPP7SL Ede1-tdimer2-aGFP + PCP-GFP(2x) | *MATa his3Δ1 leu2Δ0 met15Δ0 ura3Δ0 ADC17-24xPP7SL-LoxP EDE1-aGFP:LEU2 [pFA6:cyc1p-PCP-EGFP(2X)::HIS3]* | This study |
| Ede1-aGFP + p416-Adc17-24xPP7SL-Stop-24xMS2SL + PCP-GFP(2x) + MCP-mCherry | *MATa his3Δ1 leu2Δ0 met15Δ0 ura3Δ0 EDE1-aGFP:LEU2 [p416-Adc17-24xPP7SL-Stop-24xMS2SL + pFA6:cyc1p-PCP-EGFP(2X)-cyc1p-MCP-mCherry::HIS3]* | This study |
| adc17Δ + p416-Adc17-24xPP7SL-Stop-24xMS2SL + PCP-GFP(2x) + MCP-mCherry | *MATa his3Δ1 leu2Δ0 met15Δ0 ura3Δ0 ede1::kanMx [p416-Adc17-24xPP7SL-Stop-24xMS2SL + pFA6:cyc1p-PCP-EGFP(2X)-cyc1p-MCP-mCherry::HIS3]* | This study |
| adc17Δ Ede1-aGFP + p416-Adc17-24xPP7SL-Stop-24xMS2SL + PCP-GFP(2x) + MCP-mCherry | *MATa his3Δ1 leu2Δ0 met15Δ0 ura3Δ0 ede1::kanMx EDE1-aGFP:LEU2 [p416-Adc17-24xPP7SL-Stop-24xMS2SL + pFA6:cyc1p-PCP-EGFP(2X)-cyc1p-MCP-mCherry::HIS3]* | This study |
| Adc17-24xPP7SL Abp1-mKate2-aGFP + PCP-GFP(2x) | *MATa his3Δ1 leu2Δ0 met15Δ0 ura3Δ0 ADC17-24xPP7SL-LoxP ABP1-mKate2-aGFP:LEU2 [pFA6:cyc1p-PCP-EGFP(2X)::HIS3]* | This study |
| Adc17-24xPP7SL Abp1-mKate2 + PCP-GFP(2x) | *MATa his3Δ1 leu2Δ0 met15Δ0 ura3Δ0 ADC17-24xPP7SL-LoxP ABP1-mKate2:KanMX [pFA6:cyc1p-PCP-EGFP(2X)::HIS3]* | This study |
| Adc17-24xPP7SL Ede1-tdimer2 + PCP-GFP(2x) | *MATa his3Δ1 leu2Δ0 met15Δ0 ura3Δ0 ADC17-24xPP7SL-LoxP EDE1-tdimer2:KanMX [pFA6:cyc1p-PCP-EGFP(2X)::HIS3]* | This study |
| Adc17-24xPP7SL Sla1-mKate2 + PCP-GFP(2x) | *MATa his3Δ1 leu2Δ0 met15Δ0 ura3Δ0 ADC17-24xPP7SL-LoxP SLA1-mKate2:KanMX [pFA6:cyc1p-PCP-EGFP(2X)::HIS3]* | This study |
| Adc17-24xPP7SL Vrp1-mKate2 + PCP-GFP(2x) | *MATa his3Δ1 leu2Δ0 met15Δ0 ura3Δ0 ADC17-24xPP7SL-LoxP VRP1-mKate2:KanMX [pFA6:cyc1p-PCP-EGFP(2X)::HIS3]* | This study |
| Adc17-24xPP7SL Abp1-mKate2-aGFP + PCP-GFP(2x) ede1Δ | *MATa his3Δ1 leu2Δ0 met15Δ0 ura3Δ0 ADC17-24xPP7SL-LoxP ede1Δ::LEU2 ABP1-mKate2:KanMX [pFA6:cyc1p-PCP-EGFP(2X)::HIS3]* | This study |
| Adc17-70ntΔ (CRISPR/CAS9) | *MATa his3Δ1 leu2Δ0 met15Δ0 ura3Δ0 5'UTR-70ntΔ-ADC17* | This study |
| WT + FGH17-70ntd + Kozak | *MATa his3Δ1 leu2Δ0 met15Δ0 ura3Δ0 ADC17-24xPP7SL-LoxP ABP1-mKate2:KanMX [pFA6:cyc1p-PCP-EGFP(2X)::HIS3]* | This study |
| WT + FGH17-70nt only | *MATa his3Δ1 leu2Δ0 met15Δ0 ura3Δ0 ADC17-24xPP7SL-LoxP ABP1-mKate2:KanMX [pFA6:cyc1p-PCP-EGFP(2X)::HIS3]* | This study |
| BY4741 + FGH17-70ntΔ + Kozak | *MATa his3Δ1 leu2Δ0 met15Δ0 ura3Δ0 [p416:FGH17-70ntΔ + Kozak::URA3]* | This study |
| BY4741 + FGH17-70nt only | *MATa his3Δ1 leu2Δ0 met15Δ0 ura3Δ0 [p416:FGH17--70nt only::URA3]* | This study |
| act1-101 | *MATa his3Δ1 leu2Δ0 met15Δ0 ura3Δ0 act1-101:kanMX* | Euroscraf |
| Rpl10-GFP + FGH17 | *MATa leu2Δ0 met15Δ0 ura3Δ0 RPL10-GFP:His3MX6 [p416:FGH17::URA3]* | This study |
| Rpl10-GFP + FGH17-70ntΔ | *MATa leu2Δ0 met15Δ0 ura3Δ0 RPL10-GFP:His3MX6 [p416:FGH17-70ntΔ::URA3]* | This study |
